# Functional Mammary Gland Organoids From Human Induced Pluripotent Stem Cells

**DOI:** 10.64898/2026.10.28.762519

**Authors:** Roshni R. Nair, Maarten Ganne, Larissa Mourão, Sylvia E. Le Dévédec, Colinda L.G.J. Scheele, Peter Bouwman, Bob van de Water

## Abstract

The human mammary gland arises through sequential lineage restrictions from surface ectoderm to hormone-responsive bilayered epithelium, yet a tractable in vitro model that captures these events has remained elusive. Here, we established a stepwise differentiation protocol that directs human induced pluripotent stem cells (hiPSCs) into mammary organoids (MGOs) by sequential induction of non-neural ectoderm, mammary placodes, and lineage maturation in a defined medium. The resulting MGOs reproducibly formed polarized basal (KRT14+, TP63+) and luminal (KRT8+, GATA3+) layers, with ∼25% of luminal cells expressing oestrogen receptor (ERα). MGOs accumulated lipids, expressed β-casein upon prolactin stimulation, and retained lineage fidelity after serial passaging. Upon transplantation into cleared mouse mammary fat pads, MGOs generated branched mammary ductal networks. Single-cell transcriptomics confirmed the emergence of basal, luminal hormone-sensing, luminal secretory, and stromal populations, transcriptionally aligned with adult human breast epithelium and responsive to endocrine disruption. These findings demonstrate that hiPSCs can be reprogrammed into functional mammary epithelium, providing a platform to dissect development, endocrine signalling, and breast cancer pathogenesis.

## Introduction

The mammary gland is a defining evolutionary adaptation in mammals, enabling postnatal nourishment. Comparative developmental and fossil evidence suggests that mammary glands evolved more than 310 million years ago from ancestral skin appendages, likely modified apocrine-like sweat glands associated with hair follicles, before the evolution of nipples and the complex ductal architectures characteristic of extant mammal^1,2^. This evolutionary transition produced a highly specialized, hormone-responsive organ capable of repeated cycles of growth, differentiation, and regression across the reproductive lifespan.

Mammary gland development is initiated during embryogenesis, when epithelial placodes arise from the surface ectoderm and invades the underlying mesenchyme. Subsequent morphogenesis is staged through tightly regulated epithelial-stromal interactions governed by conserved signalling pathways, including wingless-related integration site (Wnt)/β-catenin, fibroblast growth factor (FGF), Hedgehog, and Notch signalling^3–6^. These pathways coordinate cell fate specification, ductal elongation, and branching morphogenesis, establishing distinct basal and luminal epithelial lineages. In the mature gland, basal myoepithelial cells provide structural integrity and contractile function, whereas luminal cells mediate milk synthesis and secretion. Together with populations of mammary stem and progenitor cells, these lineages confer the remarkable plasticity that enables the gland to respond dynamically to hormonal cues during puberty, pregnancy, lactation, and involution^7^.

Disruption of mammary gland development and homeostasis can arise from genetic alterations, hormonal imbalance, or environmental exposure^8–10^. Congenital abnormalities such as mammary hypoplasia or aplasia reflect perturbations of early developmental programmes^10^. Genetic disorders including Turner syndrome, characterized by partial or complete monosomy X, and Ulnar Mammary Syndrome, caused by mutations in the transcription factor TBX3, illustrate how defects in lineage specification and morphogenetic signalling can lead to absent or underdeveloped mammary tissue^11,12^. In parallel, environmental endocrine-disrupting chemicals (EDCs), including bisphenol A (BPA), phthalates, and polycyclic aromatic hydrocarbons (PAHs), may interfere with hormone signalling and developmental pathways, altering epithelial differentiation and increasing long-term susceptibility to breast cancer^10,13^. Exposure to such agents during critical developmental windows is increasingly recognized as a major determinant of lifelong mammary gland health. Breast cancer, the most common malignancy among women worldwide, exemplifies the complexity of mammary development and lineage regulation. Rather than a single disease entity, breast cancer comprises multiple molecular subtypes, including luminal A, luminal B, HER2-enriched, and basal-like or triple-negative cancers, each associated with distinct clinical behaviours and therapeutic responses^14,15^. Accumulating evidence suggests that this heterogeneity reflects perturbations in specific mammary epithelial lineages or progenitor states, linking tumour identity directly to the developmental hierarchy of the gland. Central regulators of mammary morphogenesis, such as Wnt, Notch, and FGF signalling, are frequently co-opted during tumour initiation and progression.

Despite major advances in cancer genomics and transcriptomics, the earliest events underlying breast cancer initiation remain poorly understood. Access to human foetal mammary tissue is limited, and ethical constraints preclude systematic interrogation of early developmental stages. Murine models have been indispensable for defining oncogenic signalling networks; however, fundamental interspecies differences in mammary gland architecture, developmental timing, and hormonal regulation limit their translational relevance. Conventional two-dimensional cell lines lack tissue architecture and lineage diversity, while patient-derived xenografts and tumour organoids primarily capture established disease states and are poorly suited for modelling early oncogenic events or environmental perturbations.

Three-dimensional organoid cultures derived from adult human mammary tissue have enabled important advances in studying epithelial organization, hormone responsiveness, and tumour behaviour^16–18^. However, these systems retain epigenetic and genetic histories shaped by postnatal development or disease and do not recapitulate the emergence of mammary tissue from its developmental origins. Moreover, adult tissue-derived organoids offer limited capacity to model congenital disorders, developmental exposures, or the initial consequences of tumour suppressor loss. Consequently, there remains a critical unmet need for a human model system that enables controlled investigation of mammary gland development, environmental disruption, and cancer-associated genetic alterations within a unified and physiologically relevant framework.

Human pluripotent stem cells provide a powerful opportunity to model human organogenesis in vitro, yet robust systems for mammary gland differentiation have remained elusive. Existing approaches often yield heterogeneous epithelial populations, lack appropriate lineage organization, or show limited functional maturation and endocrine responsiveness^19–21^. As a result, key questions regarding human mammary development, disease susceptibility, and cancer initiation remain inaccessible to experimental interrogation.

Here, we report the generation of functional mammary gland organoids from human induced pluripotent stem cells (hiPSCs). By directing hiPSCs through TFAP2A⁺, E-cadherin⁺, cytokeratin⁺ non-neural ectodermal progenitors with development towards LEF1^+^ and TBX3^+^ mammary placodes, we generated self-organizing organoids that recapitulated essential features of human mammary gland architecture and lineage hierarchy. These organoids contained KRT8⁺ luminal epithelial cells, including a subset of ER⁺ luminal cells, KRT14⁺ basal cells, TP63⁺, smooth muscle actin (SMA)⁺ myoepithelial cells, and a population (∼4%) of mammary stem cells and progenitor cell marked by previously reported markers including *LGR5*, *PROCR*, *ALDH1A3*, *THY1* and *CD44*^22,23^. Importantly, upon prolactin stimulation, the organoids underwent functional maturation and produced the milk protein β-casein (CSN2), demonstrating endocrine responsiveness and secretory competence. The functional capacities of mammary gland organoids were confirmed by their capacity to grow out into branched ductal networks upon transplantation into cleared mouse mammary fat pads. Using this system, we further demonstrate the utility of hiPSC-derived mammary organoids as a tractable human platform for modelling environmental endocrine disruption, which resulted in altered mammary epithelial differentiation and lineage organization. Together, these findings establish hiPSC-derived mammary organoids as a versatile and developmentally faithful model for studying human mammary gland biology, environmental pollutant and chemical hazard assessment, and early events that drive breast cancer formation.

## Results

### Stepwise induction of human mammary gland organoids

To develop human mammary gland organoids, we sought to recapitulate the sequential lineage specifications of human mammary development in vitro by guiding hiPSCs through non-neural ectoderm specification, mammary placode induction, and lineage maturation (Fig. 1 a,b, S1a). We first confirmed the pluripotency and trilineage potential of two independent hiPSC lines (SIGi001-A-23 and STBCi052-A, referred to as donor #1 and donor #2, respectively), which expressed canonical pluripotency markers and displayed expected lineage responsiveness (Fig. S1b). When dissociated to single cells and aggregated in suspension, both lines reproducibly generated spheroids that progressively increased in size and acquired epithelial organization over 13 days (Fig. 1b; Fig. S1a).

**Figure 1.**
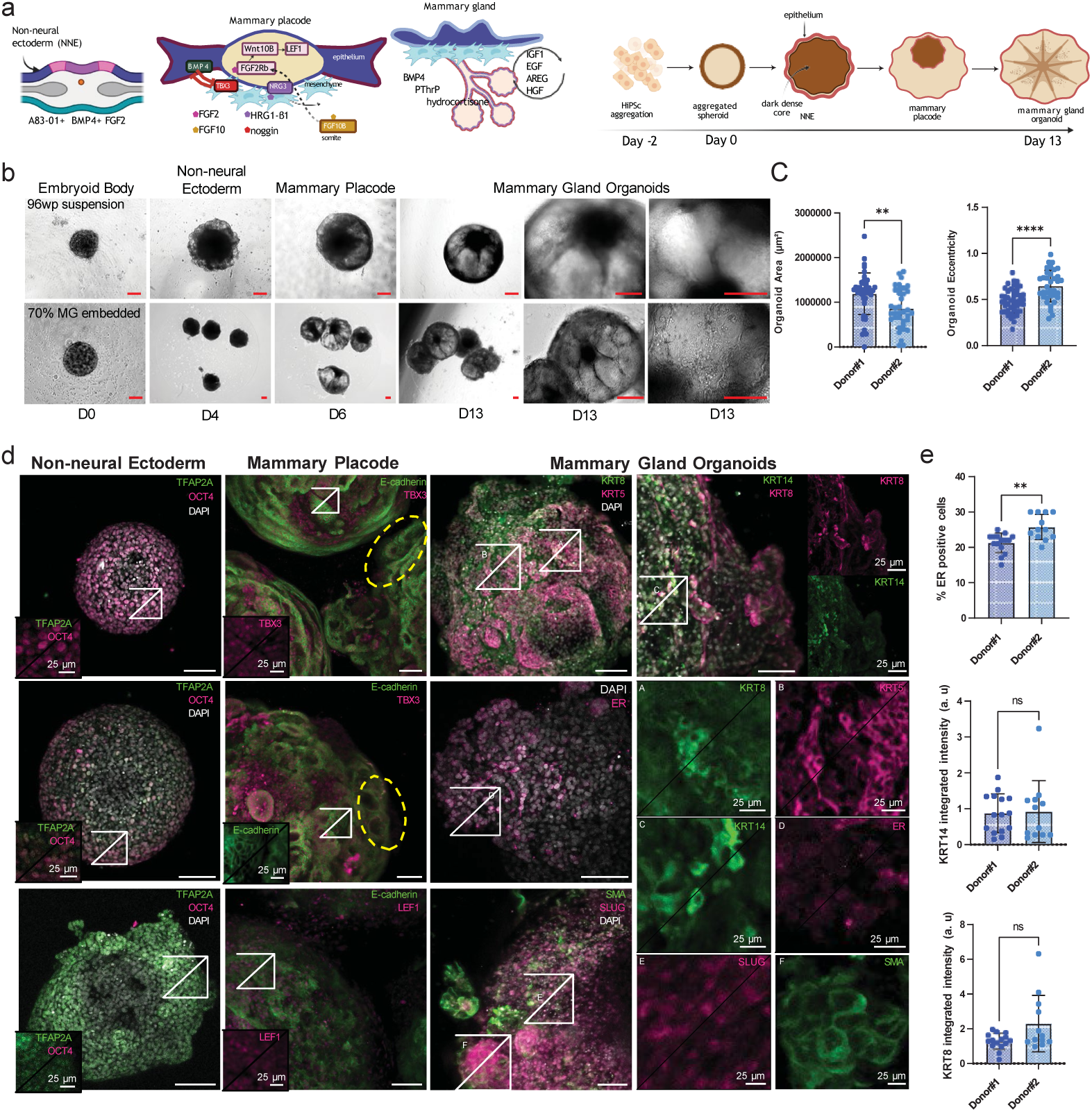
Stepwise induction of hiPSC-derived mammary gland organoids (MGOs) **a.** Schematic of mammary gland development and the corresponding in vitro differentiation strategy. Left: key stages of mammary gland development in vivo, including non-neural ectoderm (NNE), mammary placode, and mammary epithelium/mesenchyme. Right: stepwise in vitro differentiation protocol from hiPSCs. The indicated factors were added at the corresponding stages: A83-01, BMP4, and FGF2 during NNE induction; BMP4, FGF2, FGF10, noggin, and HRG1-β1 during mammary placode induction; and BMP4, PTHrP, hydrocortisone, IGF1, EGF, AREG, and HGF during mammary gland organoid differentiation. **b.** Morphological progression during mammary gland differentiation from day (D) 0 to 13. The top row shows representative organoids maintained in 96-well suspension culture, whereas the bottom row shows organoids embedded in 70% Matrigel (MG). The D13 images in the final two columns provide higher-magnification views of the organoid morphology. Scale bars, 100 μm. **c.** Morphometric analysis of mammary gland organoids from two donor lines. Brightfield images were analysed using CellProfiler and Organoid area (μm²) and eccentricity were quantified for donor #1 (n = 37 organoids) and donor #2 (n = 47 organoids). Data are shown as individual organoids, with bars indicating the mean ± s.d. Statistical significance was determined using a two-sided Mann–Whitney test; **p < 0.01 and ****p < 0.0001. **d.** Whole-mount confocal immunofluorescence characterisation of lineage progression and mammary epithelial organization. Representative images of organoids at the indicated developmental stages. Left panels: Day 4 non-neural ectoderm spheres expressed surface ectoderm markers TFAP2A and KRT8, with complete loss of OCT4. Middle panels: By day 6, placode-like structures expressed canonical placodal regulators TBX3 and LEF1 together with E-cadherin. Right panels: By day 13, MGOs acquired structures comprising KRT8+ luminal cells and KRT14+, SMA+ and SLUG+ basal/myoepithelial cells; and ER+ cells. Areas encircled by dotted yellow lines indicate descent and invagination. White boxes indicate regions shown at higher magnification in the corresponding inset panels (A–F). Insets show the cellular organisation and marker expression within these selected regions. Scale bars, 100 μm for whole-organoid images and 25 μm for magnified images. **e.**Quantification of mammary epithelial lineage-marker expression in organoids from two donor lines. The percentage of ER+ cells and integrated fluorescence intensity of KRT14 and KRT8 were quantified in donor #1 and donor #2 organoids. For ER, donor #1: n = 15 organoids; donor #2: n = 12 organoids. Data are shown as individual measurements with bars indicating the mean ± s.d. Statistical significance was assessed using a two-sided Mann–Whitney test (**p < 0.01). Integrated fluorescence intensity is reported in arbitrary units (a.u.).

To induce non-neural ectoderm, single-cell dissociated hiPSCs were seeded in U-bottom 96-well plates (10,000 cells per well) to generate uniform spheroids. Following spheroid formation, two parallel culture approaches were used: spheroids were either maintained in suspension or transferred to 70% Matrigel (MG) for continued three-dimensional culture (Fig.1b). Morphological differences were not observed in brightfield images and hence future downstream analysis were performed with suspension cultures. Spheroids were cultured in defined medium containing the TGFβ inhibitor A83-01, BMP4, and FGF2 for four days (Table S1). In this context, BMP4 promotes non-neural ectoderm differentiation while suppressing neural fates^24–26^, FGF2 enhances epithelial survival and patterning^6^, and TGFβ inhibition reinforces ectodermal commitment by blocking pro-neural cues^27^. Similar strategies have been employed to generate hiPSC-derived hair-bearing skin organoids, reflecting the evolutionary origin of the mammary gland from apocrine sweat glands^28^. By day 4, spheroids had efficiently adopted a non-neural ectodermal identity. OCT4 expression was lost (Fig. S1b), and qPCR revealed downregulation of pluripotency-associated transcripts (*OCT4*, *NANOG*) together with robust induction of *TFAP2A*, *KRT18*, and *KRT8* (Fig. S1c). Morphologically, spheroids formed compact epithelial cysts of ∼500 µm for donor 1, with smooth borders and high circularity (Fig. 1b, Fig. S1a). Across 37 organoids from donor #1 and 47 from donor #2, organoid area measured with brightfield microscopy differed significantly, with donor #1 organoids larger on average (∼1.26 × 10⁶ µm²) compared to donor #2 (∼9.61 × 10⁵ µm²). Eccentricity was higher in donor #2 than donor #1 (median ∼0.7 vs. ∼0.5) (Fig. 1c), reflecting more lobulated and branching morphologies. Immunofluorescence analysis using confocal microscopy confirmed TFAP2A and KRT8 expression at day 4 (Fig. 1d).

Between days 4 and 6, BMP inhibition combined with FGF2, FGF10, and neuregulin promoted placodal specification in the absence of direct Wnt supplementation. FGF2 and FGF10, acting through FGFR2b, are known to induce WNT10B, a marker of mammary commitment^3,5,7^, suggesting that this combination may engage an endogenous FGF–WNT signalling program to promote placodal mammary competence avoiding the need of exogenous Wnt supplementation. Consistent with activation of a WNT-responsive developmental program, LEF1 expression was increased following treatment (Fig. S1d). As a downstream effector of canonical WNT/β-catenin signalling, LEF1 provides molecular evidence that WNT-responsive transcriptional activity accompanies the acquisition of placodal identity. BMP4 and TBX3 function in complementary roles to facilitate ectodermal competence and placode specification^29^, while neuregulin further enhances TBX3 and LEF1 expression^30^. During this period, spheroids acquired placode-like features, including epithelial invagination and thickened epithelium (Fig. 1d). These morphological changes were accompanied by transient upregulation of TBX3 and LEF1 (protein expression visualized by immunofluorescence Fig. 1d; transcript levels measured by qPCR Fig. S1d), transcription factors indispensable for mammary placode initiation: *TBX3* haploinsufficiency in humans causes mammary hypoplasia^11^, while *Lef1* knockout mice lack mammary placodes^29,31^. Consistent with their embryonic roles^6^, expression peaked during placode induction and declined thereafter.

To promote maturation of the induced milk line into bilayered mammary organoids, we developed a defined medium, termed BORN (Breast Organoid Recapitulation Niche), applied over seven days. BORN medium incorporated temporally coordinated combination of paracrine and hormonal cues (Table S1) that recapitulate the mammary microenvironment. FGF2 and FGF10 to support epithelial proliferation and ductal morphogenesis^6,32^, BMP4, reintroduced at low levels, to promote epithelial stratification and luminal lineage priming; amphiregulin (AREG), an EGFR ligand transcriptionally induced by oestrogen, to mediate stromal-epithelial crosstalk required for luminal lineage commitment^33,34^; hydrocortisone and EGF acting synergistically to promote epithelial proliferation and survival^35,36^; IGF1 and HGF to enhance epithelial morphogenesis and branching^37,38^; TGFβ inhibition to facilitate epithelial polarity and structural organization^39^; PTHrP to simulate stromal feedback required for ductal outgrowth and basal cell function^40^; and oestradiol (E2) to activate oestrogen receptor signalling, essential for terminal end bud formation and amplification of AREG–EGFR loops^33,36^.

By day 13, mammary spheroids reproducibly organized into epithelial organoids (Fig. 1d, e). Immunofluorescence analysis revealed keratin-defined lineages, with presence of KRT8⁺ luminal cells and KRT5⁺/KRT14⁺ basal cells. Basal cells expressed smooth muscle actin (SMA), indicating acquisition of a myoepithelial phenotype, and co-expressed SNAI2 (SLUG), a transcription factor associated with basal lineage specification and progenitor maintenance (Fig. 1d). Within this population, subsets of luminal cells expressed oestrogen receptor (ER) indicating the emergence of hormone-responsive luminal identity^4,17,22,41–43^. Quantitative analysis showed that ∼25% of luminal cells were ER⁺ across donors (donor #1: ∼23%, n = 15 organoids; donor #2: ∼28%, n = 12 organoids) (Fig. 1e). The presence of an ER⁺ luminal population further supports the acquisition of mammary epithelial identity. The frequency of ER⁺ luminal cells were reproducible across donor lines, ranging 20-25%. KRT14 and KRT8 intensities were comparable between donors, confirming stable lineage segregation (Fig. 1e).

Bulk RNA sequencing corroborated progressive lineage restriction, separating hiPSCs, non-neural ectoderm, and mature mammary organoids. Transcriptional profiling revealed distinct basal and luminal regulatory programs that are present in the hiPSC-derived mammary gland organoids (Fig. S1e), and gene ontology analysis showed enrichment of pathways related to epithelial differentiation, branching morphogenesis, wound healing, and mammary gland development (Fig. S1f).

Together, these results demonstrate that hiPSCs can be systematically guided through conserved developmental milestones to form ER^+^ mammary organoids.

### Functional and Structural Validation of hiPSC-Derived Mammary Organoids

To evaluate the functional competency and structural fidelity of hiPSC-derived mammary organoids (MGOs), we assessed their long-term stability, secretory activation, and capacity to undergo branching morphogenesis. First we evaluated the maintenance across passages. Organoids were serially propagated by dissociation and re-aggregation across at least four passages (P0-P4) in 70% Matrigel. Within two days post-passaging, single cells extended protrusions that clustered into small colonies, which progressively self-organized into spheroids resembling early organoid formation. Brightfield imaging confirmed progressive expansion into cystic and lobulated structures (Fig. 2a). With repeated passaging in Matrigel domes beyond passage 2 (day 32+ post-P0), organoids increasingly adopted a cystic morphology, with reduced architectural complexity compared to earlier passages (P4). This shift likely reflects altered differentiation dynamics under prolonged in vitro culture. Nonetheless, immunofluorescence at passage 4 demonstrated preservation of basal cells expressing KRT14 and TP63, luminal cells expressing KRT8 and EPCAM, and subsets of luminal cells retaining ER expression (Fig. 2b). These results indicate that MGOs sustain lineage fidelity and ER+ cells despite morphological simplification over extended culture.

**Figure 2.**
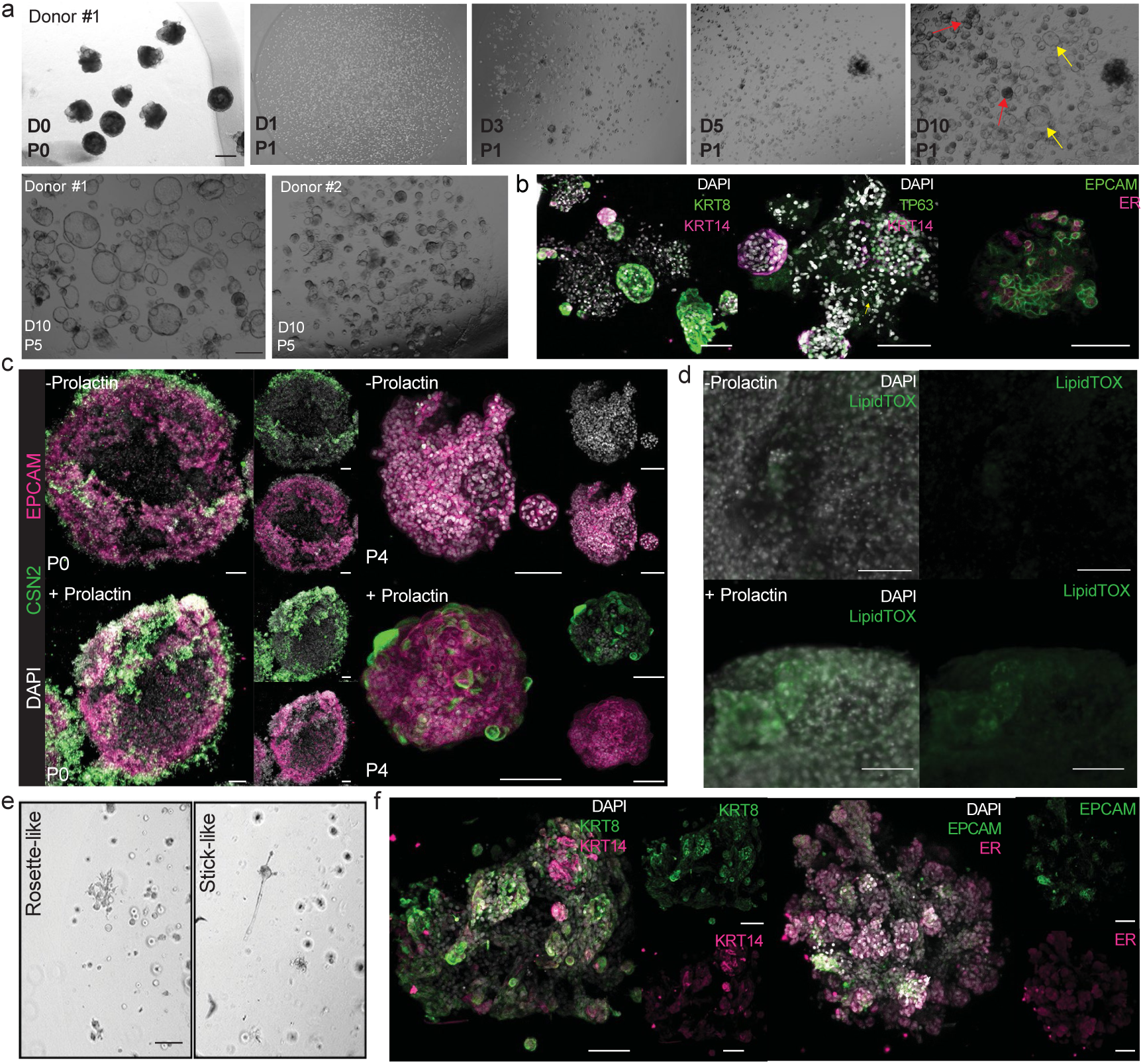
Long-term maintenance, branching morphogenesis and functional maturation of hiPSC-derived mammary gland organoids. **a.** Brightfield images of serially propagated MGOs from passage 0 (P0) to passage 5 (P5). Dissociated single cells re-aggregated to form compact (red arrow) or cystic (yellow arrow) spheroids over a period of 10 days (D0 to D10), which progressively attained cystic morphology upon further passaging, P5. Bottom panel shows P5 organoids left to right-Donor #1 and Donor #2 respectively. with passaging. Scale bars, 100 μm. **b.** Immunofluorescence confirms preservation of epithelial lineage markers during serial propagation. Immunofluorescence at P4 demonstrated continued expression of the epithelial and basal-associated markers KRT8, KRT14, TP63, supporting preservation of epithelial identity following extended culture. Scale bars, 100 μm. **c.** Prolactin-induced secretory differentiation is retained following serial propagation. Prolactin (PRL) stimulation induced β-casein (CSN2) expression in EPCAM+ luminal cells, both at early passage (P0, left) and late passage (P4, right), demonstrating retention of a hormone-responsive secretory program following extended culture. Scale bars, 100 μm. **d.** Lipid accumulation in differentiated MGOs. Brightfield (left) and confocal imaging with LipidTOX (green) confirmed abundant cytoplasmic lipid droplets in differentiated organoids, consistent with alveolar secretory maturation. Area encircled by dotted white line indicates contains droplet-resembling structures. Scale bars, 100 μm. **e.** Branching morphogenesis following mechanical fragmentation and re-embedding. Brightfield imaging of MGOs mechanically fragmented and re-embedded in Matrigel revealed distinct early branching morphologies, including rosette-like buds and elongated stick-like protrusions, demonstrating the capacity of MGOs to undergo branching morphogenesis under 3D culture conditions. Scale bars, 100 μm. **f.** Immunofluorescence demonstrates preservation of epithelial and lineage-associated marker expression in branched structures. Branched structures retained luminal KRT8-positive and basal KRT14-positive cell populations, together with ER-positive cells, consistent with preservation of epithelial markers and hormone receptor expression during branching morphogenesis. Scale bars, 100 μm.

Next, to evaluate lactogenic competence, MGOs were treated with prolactin (PRL) for 5 days. At both early (P0) and late (P4) passages, PRL induced robust expression of β-casein (CSN2) in EPCAM+ luminal cells (Fig. 2c), demonstrating that secretory capacity is maintained across passages. In parallel, functional maturation was validated by lipid accumulation, a hallmark of alveolar secretory epithelium. Brightfield imaging revealed the formation of structures resembling droplets, and LipidTOX staining confirmed abundant lipid droplets in hormone-treated organoids compared to controls (Fig. 2d). Finally, to test morphogenetic potential, lobulated MGOs were mechanically fragmented and re-embedded in 70% 3D Matrigel. Within 10–14 days, organoids generated multicellular outgrowths exhibiting two morphologies: rosette-like buds and stick-like (Fig. 2e). Immunofluorescence confirmed preservation of KRT8+ luminal and KRT14+ basal cells surrounding ER+ hormone-responsive subsets (Fig. 2f).

Together, these data show that hiPSC-derived MGOs maintain lineage fidelity across passages, show morphogenetic potential, and achieve hormone-dependent secretory activation, thereby functionally recapitulating key hallmarks of mammary gland biology.

### HiPSC-Derived Mammary Organoids Exhibit In vivo Ductal Outgrowth Capacity and Branching Morphogenesis

To assess regenerative potential in vivo, we transplanted dissociated single-cell MGO suspensions into the cleared mammary fat pads of 2.5–3-week-old immunocompromised *NOD-scid;IL2Rg^null^* (NSG) mice. Undifferentiated hiPSC spheroids and dissociated primary human breast tissue derived from reduction mammoplasty were included as controls. Successful engraftment was observed in both the MGO and primary breast epithelial groups. Whole-mount imaging revealed a spectrum of epithelial outgrowth morphologies, including branched ductal networks and compact spheroid-like structures we labelled as “*ductoids*”. Immunofluorescence confirmed the presence of basal cells expressing KRT14, and SMA, and luminal cells marked by KRT8 (Fig. 3a–c). Human-specific KU80 staining verified donor origin of the engrafted epithelium (Fig. 5c). Moreover, no outgrowths were detected from undifferentiated hiPSC spheroids (Fig. 3d,e). Importantly, ductoids from our study closely resembled those previously observed for transplanted primary human breast tissue cells ^61^, demonstrating that in vitro differentiation yields a lineage-committed mammary state capable of repopulation and morphogenesis in vivo. The fact that hiPSC-derived MGOs not only engrafted efficiently but also generated branched ductal networks at 10 weeks, underscores their enhanced morphogenetic potential.

**Figure 3.**
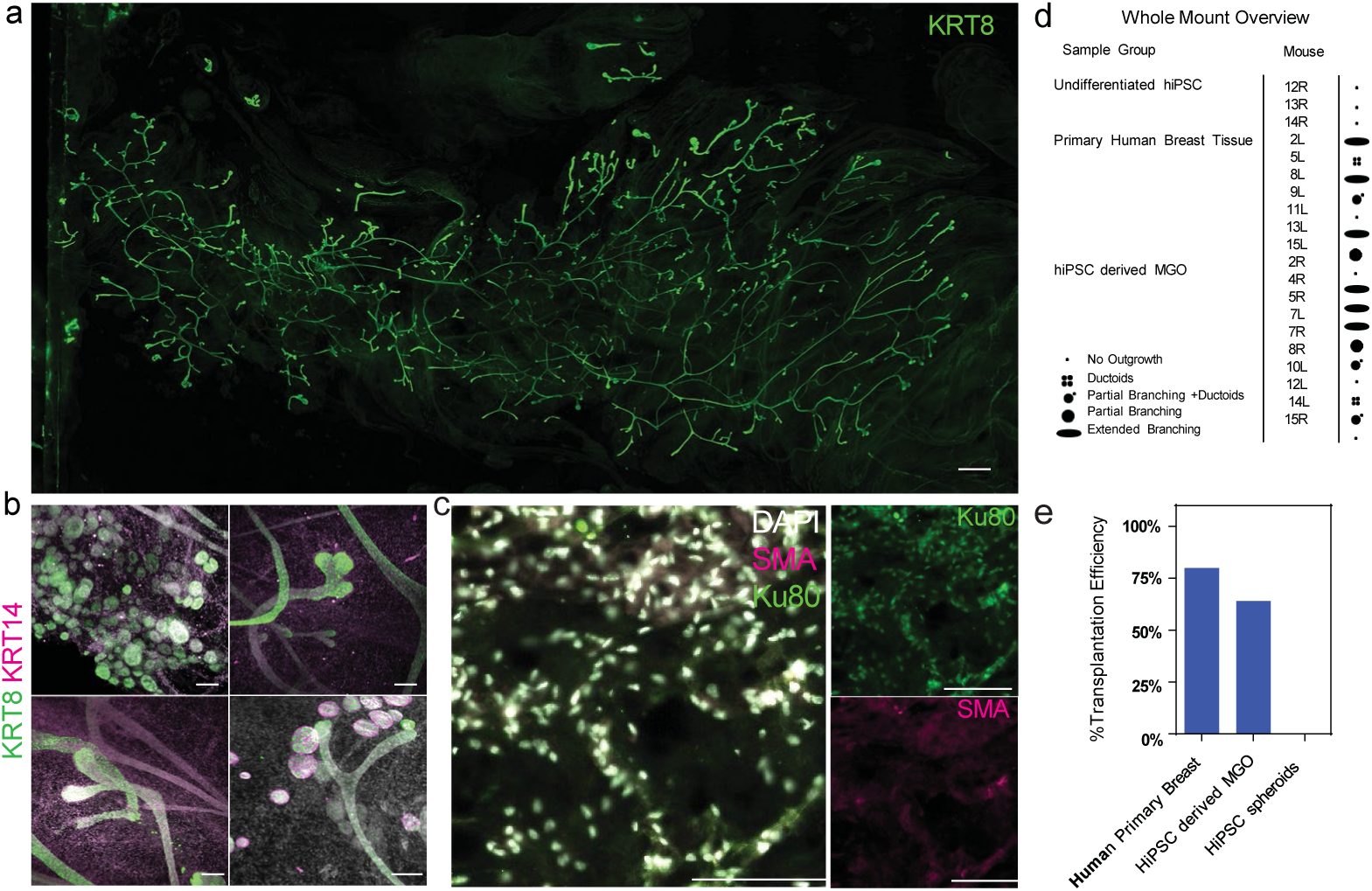
HiPSC-derived mammary gland organoids engraft and undergo ductal morphogenesis following transplantation into cleared mammary fat pads. **a.** Whole-mount analysis of epithelial outgrowth following transplantation of MGO-derived cells. Representative whole-mount image of a branched epithelial outgrowth generated following transplantation of dissociated hiPSC-derived mammary gland organoid (MGO) cells into a cleared mouse mammary fat pad. Scale bar, 1000 μm. **b.** Immunofluorescence demonstrates marker expression in branched epithelium outgrowth. Representative immunofluorescence images of regenerated epithelial structures showing bilayered organization with KRT8-positive luminal-associated and KRT14-positive basal-associated cell populations. High-resolution images showcase ductal extensions and ductoid outgrowths. Scale bars, 100 μm. **c.** Human-specific immunostaining confirms the presence of MGO-derived cells within mammary gland outgrowths. OCT imaging and immunofluorescence for the human-specific nuclear marker KU80, together with SMA, demonstrate the transplanted structures were from human MGO (n=10 glands, where 3 glands were transplanted with undifferentiated hiPSCs, 3 glands transplant-ed with cells from human breast tissue and 4 glands transplanted with MGOs). Scale bar, 100 μm. **d.** Summary table of transplantation outcomes across all whole-mount glands. Transplantation outcomes were classified according to the extent of epithelial outgrowth as no outgrowth, partial branching, ductoid formation, or extensive branching. Outcomes were classified as no outgrowth, partial branching, ductoids, or extensive branching (n=20 glands/transplants, where 3 glands were transplanted with undifferentiated hiPSCs, 7 glands transplanted with cells from human breast tissue, and 10 glands transplanted with MGOs). **e.** Quantification of transplantation efficiency across cell sources. The percentage of transplanted mammary fat pads exhibiting epithelial outgrowths was quantified following transplantation of primary human breast epithelial cells, hiPSC-derived MGOs, and undifferentiated hiPSC spheroids. The analysis includes both whole mounts and OCT sections. Bars represent the percentage of transplanted mammary fat pads exhibiting epithelial outgrowths. (n=30 glands/ transplants, where 6 glands were transplanted with undifferentiated hiPSCs, 10 glands transplanted with cells from human breast tissue, and 14 glands transplanted with MGOs).

### Single-Cell Transcriptomics of hiPSC-derived Mammary Gland Organoids Reveals Mammary Epithelial Diversity

To determine whether hiPSC-derived MGOs recapitulate the cellular diversity of the human breast epithelium, we performed single-cell RNA sequencing on day 13 organoids from suspension cultures generated from Donor #1 under baseline conditions and following 24-hour exposure to 10 µM 17 β-oestradiol (E2), 10 µM E2 plus 1 µM 4-hydroxytamoxifen (4OHT), 1 µM bisphenol A (BPA), or 10 nM diethylstilboestrol (DES) (Fig. 4a). Comparison of E2 with E2 + 4OHT enabled us the assessment of oestrogen-responsive transcriptional states, along with evaluation of oestrogen receptor-dependent responses and the antagonistic action of 4OHT. BPA and DES were further included to determine whether similar transcriptional responses could be elicited by other oestrogenic compounds and to assess the specificity of oestrogen-responsive states. Integrated analysis and unsupervised clustering of a total of 19,020 cells pooled from two biological replicates (n=10 MGOs per treatment per replicate) resolved transcriptionally distinct epithelial and mesenchymal-like populations, indicating that the differentiation protocol generated a heterogeneous mammary-like cellular ecosystem rather than a uniform epithelial spheroid (Fig. 4b).

**Figure 4.**
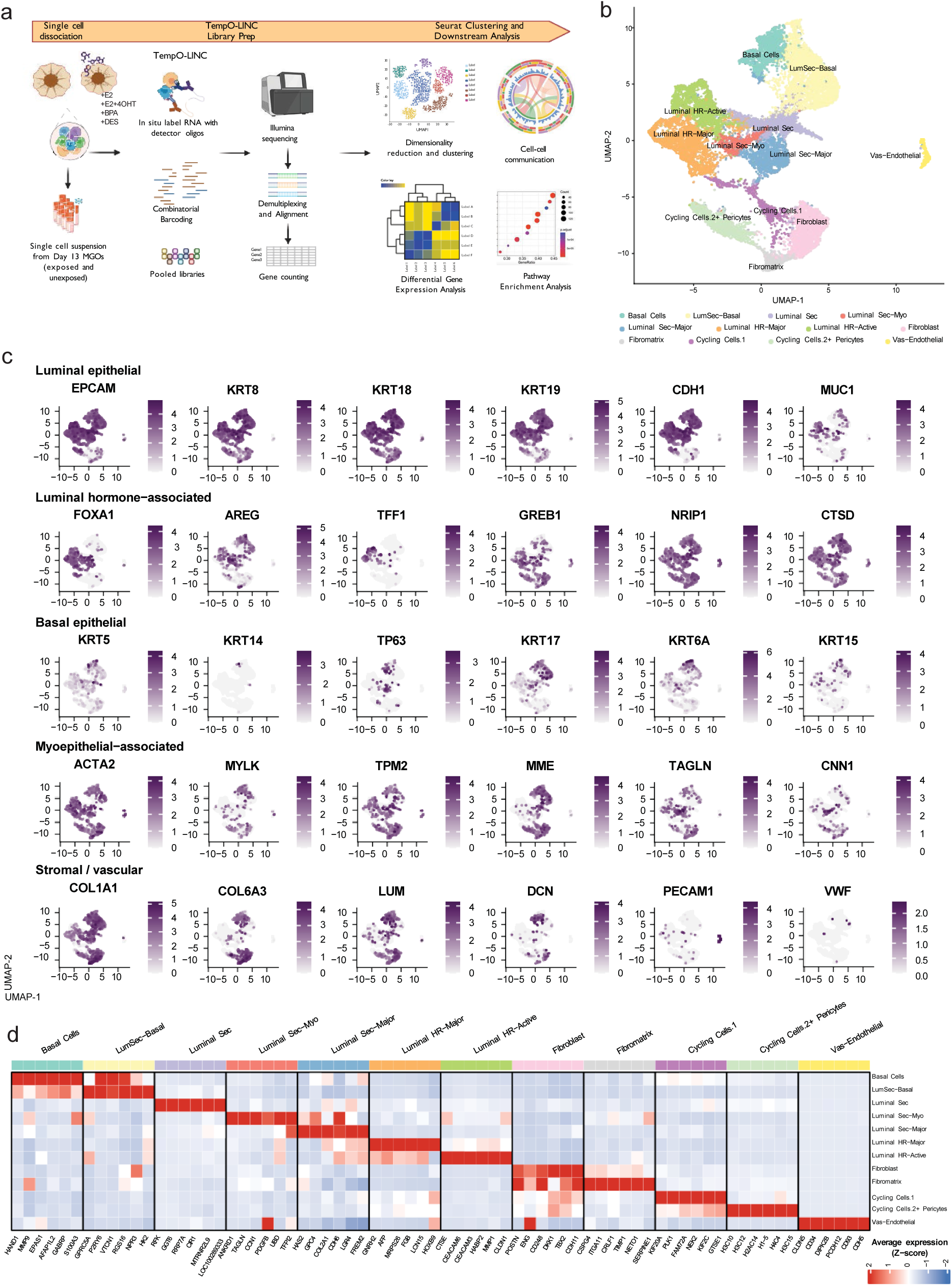
Single-cell Transcriptomics Reveals Cellular Heterogeneity and Hormonal Response in hiPSC-Derived Mammary Gland Organoids. **a.** Experimental workflow for single-cell RNA sequencing (scRNA-seq) of day 13 mammary gland organoids (MGOs). MGOs derived from donor #1 were dissociated into single-cell suspensions following exposure to the indicated endocrine conditions: untreated, 10 µM oestrogen (E2), 10 µM E2 + 1 µM 4-hydroxytamoxifen (4OHT), 1 µM bisphenol A (BPA), and 10 nM diethylstilboestrol (DES). Each condition included 10 MGOs giving approximately ∼250,000 cells/condition post fixation. Dimensionality reduction and clustering were performed on 19020 cells pooled from two biological replicates, for each replicate n=10 organoids/condition to identify transcriptionally distinct cell populations. **b.** Uniform Manifold Approximation Projection (UMAP) visualization of transcriptionally distinct cell populations in day 13 MGOs. **c.** UMAP feature plots showing expression of representative lineage and cell-state markers. Expression intensities are displayed using gene-specific colour scales. **d.** Heatmap of top 6 marker identified in each cluster defining the cell populations. The heatmap shows representative differentially expressed marker genes associated with each transcriptionally defined cell population. Colours represent scaled average gene expression (Z-score), with positive values in red indicating higher relative expression and negative values in blue indicating lower relative expression across the cell populations.

Cluster annotation identified luminal epithelial populations expressing *EPCAM*, *KRT8*, *KRT18*, *KRT19* and *CDH1*, alongside luminal hormone-responsive populations characterized by *FOXA1* and the oestrogen-associated genes *AREG*, *TFF1*, *GREB1* and *NRIP1* (Fig. 4c). Basal-associated populations expressed *KRT5*, *KRT14*, *TP63*, *KRT17* and *KRT6A*, whereas additional expression of *ACTA2*, *MYLK*, *TPM2*, *MME* and *TAGLN* supported the presence of myoepithelial-associated transcriptional features (Fig. 4c). Additional luminal differentiation and oestrogen-associated markers, including *SERPINA1*, *TFF3*, *CTSD* and *AGR2*, were identified in the transcriptional data (Table S2)^23,44–47^. Together, these expression patterns broadly correspond to the basal/myoepithelial, luminal secretory and luminal hormone-responsive compartments described in adult human breast single-cell atlases.

In addition cells expressing the progenitor- and stem-associated markers SOX9, ALDH1A3, LGR5 and PROCR were also detected within the organoid cultures (Fig. S2a), consistent with the presence of transcriptional features associated with mammary progenitor-like states^22,48^. However, expression of these markers alone does not establish functional stem-cell identity. In addition to epithelial populations, stromal-like cells expressed extracellular matrix-associated markers, including *COL1A1*, *COL6A3*, *LUM* and *DCN*, while *PECAM1* expression supported the presence of a vascular-associated population (Fig. 4c). Together, these patterns indicated that the organoids contained multiple mammary epithelial states alongside stromal- and vascular-associated transcriptional populations.

We next examined the transcriptional programs underlying the annotated populations to determine the extent to which these broad identities corresponded to established mammary cell states (Fig. 4d). Basal and basal–luminal populations showed enrichment of basal/myoepithelial-associated genes, including *HAND1, MMP9, EPAS1* and *GABRP*, while a distinct myo-epithelial population expressed *ANKRD1, TAGLN, CCN1* and *PDGFB*, consistent with contractile and extracellular-matrix-associated differentiation. The luminal compartment was similarly heterogeneous, with a secretory population expressing *HAS2, LGR4* and *FREM2*, alongside distinct hormone-responsive states. Fibroblast-like cells were characterized by expression of markers like *POSTN* and *CDH11*, whereas endothelial cells expressed canonical vascular markers including *CLDN5* and *CDH5*. A distinct proliferative population was marked by expression of mitotic genes, e.g., *KIF20A, PLK1*, and *NEK2* (Fig. 4d).

Cell-cycle scoring based on S-phase and G2/M-associated gene expression demonstrated that this ‘Cycling Cells 1’ population exhibited markedly higher S and G2/M score values compared with the majority of differentiated epithelial and stromal populations (Fig. S2b). Elevated cell-cycle scores were also observed in the ‘Cycling Cells 2 + Pericytes’ population, suggesting the presence of proliferative subpopulations within these compartments. In contrast, most luminal, basal, fibroblast, and endothelial populations showed relatively low S and G2/M score values, consistent with a predominantly non-cycling state. The putative pericyte population was dominated by histone transcripts among its top-ranked genes and showed relatively limited enrichment for canonical mural-cell-associated genes, such as *RGS5*, *PDGFRB*, *CSPG4*, *MCAM*, and *KCNJ8* (Table S2). Thus, the transcriptional evidence supporting a pericyte identity was comparatively weak. Collectively, these profiles support broad mammary epithelial and stromal identities while indicating that several epithelial populations retain features of intermediate or developmental states.

Gene Ontology analysis showed enrichment for mammary gland development, mammary epithelial morphogenesis, oestrogen receptor signalling, steroid hormone response and branching morphogenesis pathways (Fig. S2c). Thus, the organoids did not merely express isolated mammary markers but organized into transcriptionally defined cell states associated with mammary epithelial development and endocrine function. Importantly, non-mammary lineages such as skin keratinocytes (*KRT1*, *MLANA*) or neuroectodermal derivatives (*PAX6*, *NRXN1*, *DLX1*) were absent or minimal (Fig. S2d), supporting the specificity of directed differentiation. Together, these data indicate that hiPSC-derived MGOs acquire multiple mammary lineage-associated transcriptional states rather than a uniform epithelial identity. While several populations, particularly fibroblast-like, endothelial and cycling cells, display coherent lineage-associated programs, the epithelial populations show greater transcriptional heterogeneity. The MGOs therefore capture broad cellular diversity of the human mammary tissue while retaining features of transitional or developmentally immature states.

To directly benchmark organoid-derived cell identities against primary human breast tissue, we performed label transfer using the Kumar et al. adult human breast atlas as a reference^46^. Organoid-derived cells mapped onto luminal hormone-responsive, luminal secretory, basal, fibroblast, endothelial and pericyte-associated populations, supporting transcriptional alignment with the cellular architecture of adult human breast tissue, with a mean prediction confidence of 69.1% across broad cell-type assignments (Fig. S2e). The strongest concordance was observed for epithelial compartments, whereas immune and mature vascular populations were underrepresented, consistent with the expected limitations of a predominantly epithelial organoid system lacking immune and vascular niche inputs.

### Hormone Responsive Cell States in hiPSC-Derived Mammary Gland Organoids

To gain insight into the potential application of our MGOs for oestrogenic hazard assessment, we investigated how the transcriptionally defined organoid cell states varied across endocrine exposure conditions. Comparison of the relative proportions of annotated cell populations revealed differences in cellular composition among untreated MGOs and organoids exposed to E2, E2 + 4OHT, BPA or DES. In particular, BPA- and DES-exposed organoids exhibited a lower relative representation of Lum Sec-Basal and Lum HR cells, and a higher representation of Lum Sec and stromal-associated and cycling populations compared with untreated MGOs.(Fig. 5a).

**Figure 5.**
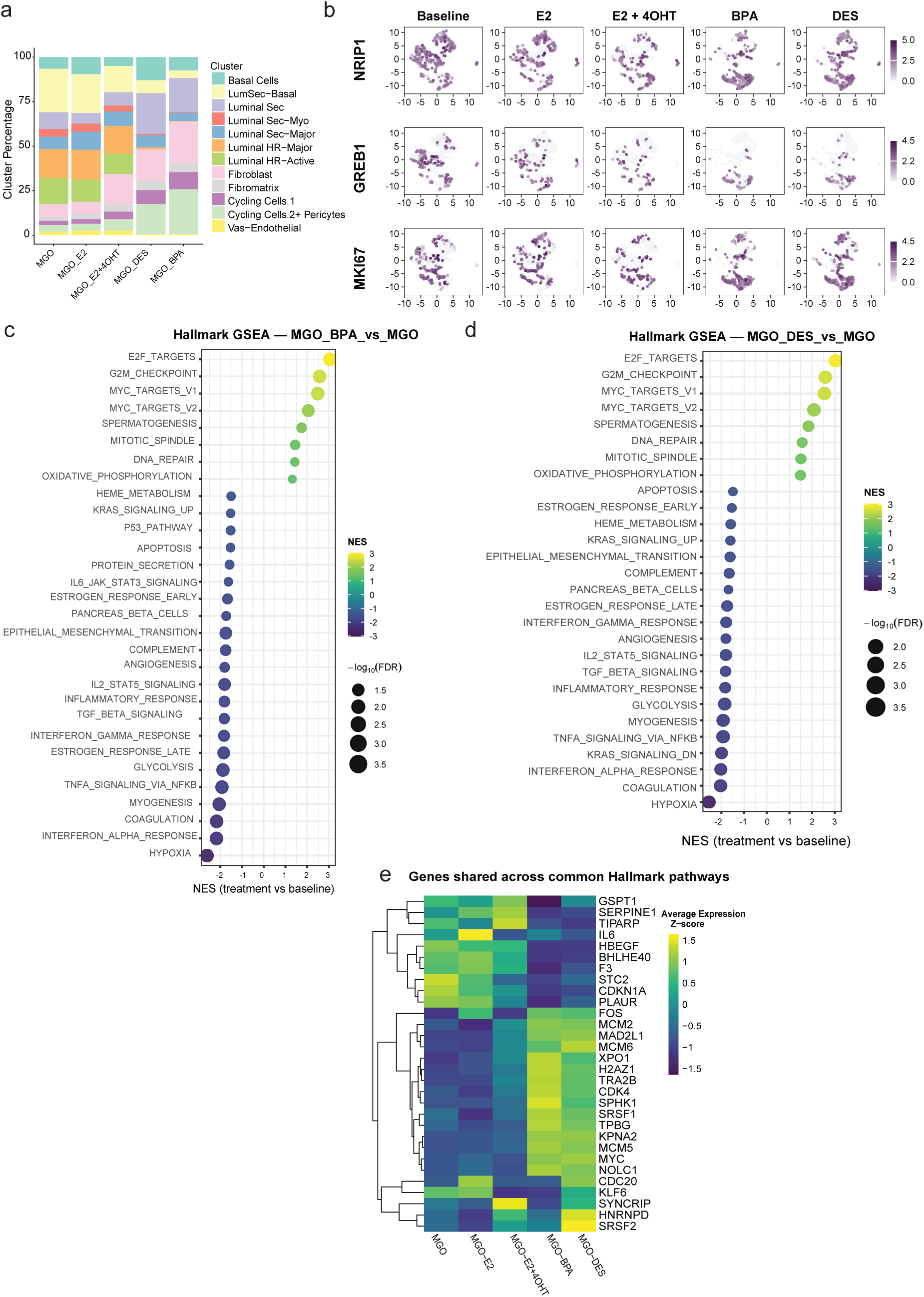
Endocrine perturbation remodels hormone-responsive transcriptional programs in hiPSC-derived mammary gland organoids. **a.** Relative abundance of individual cell populations across untreated (MGO) and endocrine-treated organoids (MGO_E2, MGO_E2+4OHT, MGO_DES and MGO_BPA), demonstrating treatment-dependent alterations in cellular composition. **b.** UMAP projections showing the expression of NRIP1, GREB1, and MKI67 across the five experimental conditions. Cells are displayed in a shared UMAP embedding, with colour intensity indicating normalized gene expression. **c.** Hallmark GSEA comparing 1 µM BPA-treated MGOs with untreated MGOs. d. Hallmark GSEA comparing 10 nM DES-treated MGOs with untreated MGOs. **e.** Heatmap showing the expression of genes shared across enriched Hallmark gene sets in mammary gland organoids (MGOs). Hierarchical clustering reveals distinct expression patterns across untreated MGO, E2-, E2 + 4OHT-, BPA- and DES-treated organoids. Colours indicate gene-wise Z-scores of average RNA expression across conditions.

Given the differences in the relative representation of hormone-responsive luminal and cycling cell populations, we next examined the distribution of selected hormone-associated and proliferation-associated transcripts across exposure conditions. We visualized *GREB1*, an established oestrogen-responsive gene, *NRIP1*, a nuclear receptor-associated transcriptional coregulator, and *MKI67*, a marker of cell proliferation, on condition-specific UMAPs (Fig. 5b). In line with previous observations in human^49^ and mouse^50^ mammary gland tissue, *NRIP1* expression was observed beyond the luminal epithelial populations, including regions of the UMAP occupied by fibroblast-like and cycling cell populations. *GREB1* and *MKI67* also displayed heterogeneous expression patterns across the exposure conditions. Notably, upon treatment with 1 µM BPA or 10 nM DES the distribution of cells expressing *NRIP1*, *GREB1*, or *MKI67* shifted towards stromal and cycling populations, suggesting lineage plasticity (Fig. 5b).

To examine relationships among epithelial states, we performed trajectory inference across the epithelial compartment. Trajectory inference across epithelial populations identified a continuous differentiation axis spanning basal, luminal secretory and luminal hormone-responsive states (Fig. S3a–d, Tables S3, S4). The inferred trajectory described a continuous transcriptional axis spanning hormone-responsive luminal, secretory luminal, basal-associated and myoepithelial-like states rather than a simple linear differentiation hierarchy. Lum HR cells occupied early positions along the inferred trajectory, followed by distinct Lum Sec subpopulations and basal/myoepithelial-associated states, with a further Lum Sec population occupying later positions. Because pseudotime represents the relative organisation of transcriptional states rather than chronological developmental time, these relationships do not establish a developmental sequence or lineage relationship between the identified populations. Instead, the trajectory captures continuous transcriptional variation across epithelial states. Hormonal receptor modulation significantly redistributed cells along this epithelial trajectory (Kruskal–Wallis p < 2.2 × 10^-16^). Untreated MGOs occupied the earliest median pseudotime state, while E2 and E2+4OHT produced modest but significant shifts. In contrast, BPA and DES caused larger displacements along the pseudotime, with 1 µM BPA showing the strongest shift among the tested perturbations, where median pseudotime increased from 0.039 in untreated organoids to 0.247 following BPA exposure and 0.128 following 10 nM DES exposure. These shifts are consistent with the altered epithelial-state composition observed in Fig. 5a, although pseudotime differences may reflect changes in cell-state abundance, transcriptional state, or both. Thus, the trajectory analysis provides evidence that 24-hour endocrine perturbations altered the organisation of epithelial transcriptional states.

To define the molecular programs associated with epithelial trajectory progression, we identified genes showing significant spatial autocorrelation along the Monocle3 trajectory (q < 0.05, Moran’s I > 0.1) and examined their expression patterns across pseudotime. (Table S5). The early pseudotime regions were characterized by expression of developmental and epithelial markers including *IGF2, DLK1, CD24* and *GOLM1*, whereas later states were enriched for genes associated with epithelial remodelling and secretory differentiation, including *TGFBI, GABRP, HAND1, IGFBP3, MUC16* and *NPR3* (Fig. S3e). FGSEA Gene set enrichment analysis further revealed significant associations with oestrogen-response, glycolysis, hypoxia, inflammatory signalling, epithelial-mesenchymal transition and PI3K-AKT-mTOR pathways, indicating remodelling of developmental, metabolic and endocrine programs across the epithelial trajectory (Fig. S3f). Together, these analyses demonstrate that MGOs contain dynamic epithelial state transitions that remain responsive to hormonal and environmental perturbations.

We next investigated how modulation of oestrogen signalling affected the transcriptional state of the organoids. Nine Hallmark signatures were significantly enriched (FDR-adjusted P < 0.05) across three comparisons: E2+4OHT vs E2, BPA vs MGO, and DES vs MGO (Table S7). Notably, the oestrogen response early signature was negatively enriched in the E2 + 4OHT versus E2 comparison (NES = −1.57), indicating antagonistic effect of 4OHT. This was accompanied by enrichment of cell-cycle-associated Hallmark pathways, including E2F targets, MYC targets and G2/M checkpoint signatures (Table S6,7). An additional 18 Hallmark signatures were shared between BPA and DES but did not reach the significance threshold in the E2 + 4OHT comparison ( Table S7). These included positively enriched DNA repair, mitotic spindle and MYC targets V2 signatures, as well as negatively enriched epithelial–mesenchymal transition, TGF-β signalling, oestrogen response late and immune-associated signatures (Fig. 5c,d, Table S6,7). These responses are consistent with prior observations that endocrine-disrupting compounds can perturb mammary epithelial architecture and signalling^51–54^., but here extend those findings to a developmentally derived, hiPSC-based human mammary model.

To examine the expression patterns of genes associated with the nine Hallmark signatures shared across all three comparisons, we selected genes represented in at least three positively enriched or three negatively enriched Hallmark gene sets. Gene-wise standardized average expression analysis across the five conditions revealed differences in the relative expression of these pathway-associated genes (Fig. 5e). Several cell-cycle- and DNA-replication-associated genes, including *MYC* and established MYC target genes like *MCM2*, *MCM5*, *MCM6*, *CDK4*, and *MAD2L1*, ^55^, showed higher relative expression in BPA- and DES-exposed organoids. Conversely, the relative expression of *CDKN1A* and *PLAUR* was decreased upon BPA or DES exposure, in line with the fact that these genes are repressed by *MYC* ^56,57^. These gene-level patterns suggested that induction of *MYC* plays a key role in the response of MGOs to 1 µM BPA or 10 nM DES.

To investigate whether the cellular diversity observed within MGOs was accompanied by coordinated multicellular signalling, we performed ligand-receptor interaction analysis across all identified epithelial and stromal populations (Fig. S4a). Extensive communication was predicted between basal, luminal secretory, luminal hormone-responsive, fibroblast/fibromatrix, endothelial and pericyte-associated compartments, suggesting a highly interconnected signalling network within the organoid niche. Communication was not restricted to epithelial populations but instead involved widespread reciprocal interactions between epithelial and stromal cell states, consistent with the epithelial-stromal crosstalk that underpins mammary gland development and tissue homeostasis.

Network-level analysis identified fibromatrix cells as a dominant signalling hub within the organoid ecosystem. Fibromatrix populations exhibited the greatest interaction burden and engaged in extensive bidirectional communication with basal, luminal secretory and hormone-responsive epithelial compartments (Fig. S4a, b; Tables S8a-i). Basal cells similarly displayed high signalling activity, particularly through reciprocal interactions with fibromatrix populations, consistent with the known association between basal epithelial cells and extracellular matrix-rich stromal compartments in the mammary gland. Fibroblast and endothelial populations further contributed to the signalling landscape through extensive interactions with epithelial cell states, indicating the emergence of a multicellular microenvironment rather than an isolated epithelial culture.

Examination of the highest-confidence ligand–receptor interactions revealed several signalling pathways previously implicated in mammary morphogenesis and epithelial maintenance. Prominent extracellular matrix-associated interactions included FN1–CD44, FN1–SDC1, FN1–SDC2 and FN1–PLAUR signalling, linking epithelial and stromal populations (Fig. S4b). Growth factor signalling was also highly represented, including IGF2–IGF1R, IGF2–INSR and IGF2–IGF2R interactions, as well as VEGFA-mediated communication with endothelial-associated populations. In addition, multiple interactions involved NOTCH-family receptors, including DLK1–NOTCH1/2/3/4 and APP–NOTCH2 signalling, supporting the presence of developmental signalling networks associated with lineage specification and epithelial differentiation. Collectively, these findings suggest that MGOs establish signalling programs characteristic of a developing mammary niche and not merely a collection of mammary-like cell types.

Our single-cell sequencing data demonstrate that hiPSC-derived MGOs contain major epithelial states of the human mammary gland, including transcriptionally hormone-responsive luminal populations, and that these populations respond dynamically to endocrine and environmental perturbations.

## Discussion

Here, we establish a stepwise strategy for generating human mammary gland organoids (MGOs) from hiPSCs by recapitulating key developmental transitions of mammary gland formation *in vitro*. By sequentially directing hiPSCs through non-neural ectoderm specification, mammary placode induction and epithelial maturation, we generated organoids containing distinct basal and luminal compartments, including an ER⁺ hormone-responsive population. The resulting MGOs displayed functional mammary characteristics, including prolactin-induced β-casein expression and lipid accumulation, and demonstrated the capacity to generate human ductal outgrowths following transplantation. Single-cell transcriptomic analysis further revealed diverse mammary epithelial and stromal populations and demonstrated that these states remain responsive to hormonal and environmental endocrine perturbation. Together, these findings establish MGOs as a developmentally derived human model for investigating mammary epithelial differentiation, endocrine signalling and tissue morphogenesis.

Our differentiation approach was designed to mimic major developmental transitions occurring during mammary gland specification, progressing through non-neural ectoderm, mammary placode formation and epithelial lineage maturation. The resulting organoids reproducibly generated epithelial structures composed of KRT14⁺ basal and KRT8⁺ luminal compartments, including ER⁺/GATA3⁺ hormone-responsive luminal cells. Importantly, maintenance of ER^+^ epithelial populations remains a persistent challenge in mammary culture systems, making the presence of a stable hormone-responsive compartment a notable feature of the model. This is relevant not only for basic research into hormonal regulation of human mammary gland development, but also for studying the impact of oestrogenic mammary gland carcinogens. While prolactin-induced milk protein expression supported mammary gland specific differentiation, we did not assess in vitro branching capacity of MGOs beyond the formation of multicellular outgrowths. Whether these multicellular outgrowths are indicative of true branching potential remains to be determined. This may require long term culturing, as has previously been shown for multilineage mouse embryonic stem cell-derived mammary organoids^58^, or adaptations such as alternating FGF2 and epidermal growth factor (EGF) treatments to induce branching and elongation ^59^.

Nevertheless, transplantation into cleared fat pads demonstrated that hiPSC-derived cells can engraft and form ductal outgrowths *in vivo*, establishing structural fidelity with bilayered organization. In contrast to previously published protocols^19–21^, our MGOs did not require the addition of human fibroblasts to support ductal outgrowth *in vivo*. This may be attributed to the fact that fibroblast-like cells and progenitors are already present in our cultures. It is also possible that our MGO differentiation protocol yielded more mammary gland stem cells than previously published methods. Similarly, our primary breast tissue control samples grew out into mammary gland ductal networks without a need for co-transplanted human fibroblasts^60^, most likely because they were enriched for regenerative capacity^61^. Besides ductal networks, we also observed what we termed ‘ductoids’: compact spheroids or cysts that resemble structures previously seen upon transplantation of human ^62^ or bovine ^63^ primary mammary gland cells. The formation of these structures may reflect suboptimal experimental conditions pre or post transplantation, but taken together, the *in vivo* outgrowths confirm the *in vitro* characterization of MGOs as a model for human mammary gland development.

The formation of MGO-derived human mammary gland networks *in vivo* provides a potential means to study (pre)neoplastic changes that are important for cancer initiation and promotion. Proof of principle that mutant hiPSC-derived mammary gland cells can grow out into tumours in mice has been shown by Liu et al. ^21^ using subcutaneous and orthotopic transplantations. If cancer initiation can also occur upon mammary gland reconstitution, this would allow studying early tumour development in a physiologically more relevant context. Moreover, this may provide insights in how to recapitulate critical steps in cultured MGOs. Combining defined endocrine exposures with genetic perturbation and transplantation could further enable investigation of whether exposure-associated epithelial states influence susceptibility to subsequent oncogenic alterations.

Single-cell analysis further places these observations within a broader cellular context. The organoids contain multiple epithelial states corresponding to basal, luminal secretory and hormone-responsive populations, together with stromal-associated and cycling populations, all aligning with the cellular hierarchy of the human breast. Crucially, non-mammary lineages (e.g., neuroectoderm, skin keratinocytes) were absent, underscoring the specificity of directed differentiation. Cell-cycle scoring further revealed proliferative heterogeneity within the organoids. S-phase and G2/M-associated transcriptional programs were strongly enriched in defined cycling populations, whereas most differentiated epithelial and stromal populations exhibited comparatively low cell-cycle scores. These findings indicate that the iPSC-derived breast organoids comprise both proliferative and more differentiated cellular states rather than representing a static cellular population.

The inferred epithelial continuum further supports a model in which mammary epithelial identity is organised as a dynamic spectrum rather than a set of completely discrete populations. Changes in the distribution of cells along this transcriptional landscape following endocrine exposure suggest that these states remain responsive to environmental signals. Trajectory analysis revealed a continuous epithelial continuum spanning basal, luminal secretory and hormone-responsive states, while endocrine perturbations significantly altered cell-state distributions along this trajectory. These findings suggest that hormone exposure influences mammary epithelial organization not only through direct transcriptional regulation but also through modulation of epithelial state dynamics.

The ability to investigate endocrine-associated transcriptional responses represents an important strength of this platform. Treatment with the established oestrogenics BPA and DES exhibited overlapping changes in epithelial-state representation and transcriptional programs. In particular, both compounds were associated with reduced representation of hormone-responsive luminal cells, increased representation of secretory luminal cells and shifts in epithelial pseudotime distributions. At the pathway level, BPA and DES shared positive enrichment of cell-cycle-associated signatures, including E2F targets, G2/M checkpoint and MYC targets, alongside negative enrichment of oestrogen-response and several signalling-associated signatures. Together, these observations raise the possibility that endocrine-active compounds influence the balance between mammary epithelial differentiation and proliferation-associated transcriptional states. Although these findings do not establish direct lineage transitions or altered proliferative activity, they highlight the potential of MGOs to investigate how endocrine exposure is associated with changes in mammary epithelial organization.

The predicted ligand–receptor interactions suggest that these responses occur within a multicellular context rather than in isolation. The presence of fibroblast-like, endothelial and other stromal-associated populations provides a potential source of paracrine and extracellular-matrix signals that may contribute to epithelial organisation and endocrine responsiveness. Although these computationally inferred interactions require experimental validation, they point towards a broader advantage of the organoid system: the possibility of studying epithelial–niche interactions alongside epithelial differentiation and hormone response within the same human model.

Despite these advances, limitations remain. Long-term passaging over 6-8 weeks reduces architectural complexity, suggesting that niche signals are critical to sustain morphogenetic capacity. In vivo, mammary epithelial cells reside in a highly specialized microenvironment, comprising fibroblasts, adipocytes, vasculature, immune cells, and extracellular matrix^46^. MGOs, while bilayered and hormone-responsive, currently lack this multicellular niche. Future integration of stromal and vascular components, perhaps through co-culture or bioengineered scaffolds^64–66^, will be essential to recapitulate ductal remodelling, alveologenesis, and full lactation. The endocrine-exposure findings should be interpreted as exploratory and require further analysis using concentration ranges to determine physiologically relevant transcriptional responses. Furthermore, changes in pseudotime distributions and whole-organoid pathway enrichment may reflect differences in cell-state composition rather than direct transcriptional regulation or lineage transitions. Independent biological replicates and cell-type-resolved functional studies will be needed to establish the reproducibility and biological consequences of these observations.

Overall, the value of this model lies in the integration of developmental specification, epithelial heterogeneity, functional hormone responsiveness and tissue-level morphogenesis within a genetically tractable human system. In particular, the presence of various distinct mammary gland (progenitor) populations, including ER⁺ luminal states, provides an opportunity to investigate how mammary epithelial states are established and subsequently remodelled by hormonal and environmental signals. This framework should be useful not only for studying human mammary development, but also for defining cellular responses to endocrine perturbation and, ultimately, for modelling disease-associated changes in a human genetic background.

## Acknowledgements

We are grateful to Dr. Wim De Maerteleire (Mediclinic, Belgium) for providing the primary human breast tissue and Pauline Luo (VIB–KU Leuven Center for Cancer Biology) for technical assistance with cell culture and primary tissue isolation for the transplantation studies. We thank Kostas K. Tassis (Leiden Academic Centre for Drug Research, Leiden University) for microscopy support. We also acknowledge the Cell Observatory, Leiden University, which is part of NL-BioImaging and Euro-BioImaging, for imaging support.

Figures were assembled using Adobe Illustrator, and selected scientific illustrations were created using BioRender.com under a licensed subscription.

This work was carried out within the framework of the European Partnership for the Assessment of Risks from Chemicals (PARC) and received funding from the European Union’s Horizon Europe Research and Innovation Programme under Grant Agreement No. 101057014. The mouse work was carried out under Stichting Tegen Kanker (STK) funding under mandate numbers 2023-045. The views and opinions expressed are those of the authors only and do not necessarily reflect those of the European Union or the European Health and Digital Executive Agency (HaDEA) or STK. Neither the European Union nor the granting authority can be held responsible for them.

## Methods

### Human pluripotent stem cell lines and culture

Experiments were performed using two independent human induced pluripotent stem cell (hiPSC) lines derived from healthy female donors: SIGi001-A-23 (passages 58–65; https://hpscreg.eu/cell-line/SIGi001-A-23, donor #1) and STBCi052-A (passages 16–23; https://hpscreg.eu/cell-line/STBCi052-A, donor #2). Both cell lines tested negative for mycoplasma contamination prior to use.

Cells were maintained on growth factor–reduced Matrigel-coated plates (0.144 mg/mL, lot-matched) in mTeSR1 medium (STEMCELL Technologies), supplemented with penicillin–streptomycin. Medium was refreshed daily until cultures reached approximately 70–80% confluency, at which point cells were passaged every 4–5 days using ReLeSR according to the manufacturer’s instructions, yielding small cell clusters. For single-cell dissociation, cultures were treated with Accutase and replated in mTeSR1 supplemented with 10 μM Y-27632 dihydrochloride (ROCK inhibitor) to enhance cell survival. Twenty-four hours after seeding, medium was replaced with mTeSR1 without Y-27632. Cells were maintained at 37 °C in a humidified incubator with 5% CO₂.

### Differentiation of hiPSCs into mammary gland organoids

hiPSC colonies at approximately 70–80% confluency were dissociated into single cells using Accutase for 5 min, washed, and seeded at 10,000 cells per well in U-bottom ultra–low-attachment 96-well plates in mTeSR1 medium supplemented with penicillin–streptomycin and 10 μM Y-27632 dihydrochloride. Cell aggregation was facilitated by centrifugation at 300g for 5 min. Cultures were incubated at 37 °C with 5% CO₂ for 48 h to generate embryoid bodies (day −2 to day 0).

On day 0, spheroids were transferred to surface ectoderm induction medium consisting of Advanced DMEM/F12++ supplemented with B27, 1 μM A83-01, 20 ng/mL FGF2, 10 ng/mL BMP4, and 2% Matrigel. Medium was refreshed every 48 h by partial exchange (approximately 85–90 μL per well). After 4 days, aggregates exhibited cyst-like morphology and marker expression consistent with non-neural ectoderm identity.

Placode induction was initiated on days 4–5 by culturing spheroids in Advanced DMEM/F12++ supplemented with B27, 100 ng/mL Noggin, 50 ng/mL FGF2, 25 ng/mL FGF10, 25 ng/mL neuregulin, and 2% Matrigel. From day 6 onward, aggregates were cultured in mammary organoid induction medium (BORN medium), consisting of Advanced DMEM/F12++ supplemented with B27, 50 ng/mL FGF2, 5 ng/mL BMP4, 50 ng/mL FGF10, 40 ng/mL neuregulin, 10 ng/mL HGF, 25 ng/mL amphiregulin, 25 ng/mL EGF, 20 ng/mL IGF1, 1 μg/mL hydrocortisone, 0.2 nM 17β-oestradiol, 50 ng/mL parathyroid hormone–related protein (PTHrP), and 5% Matrigel. BORN medium was refreshed every 2 days. Organoids were maintained under these conditions until day 13, at which point they exhibited a bilayered architecture comprising luminal and basal epithelial lineages. All cytokines and growth factors used were recombinant human proteins. Complete medium compositions and reagent sources are provided in Table S1.

### Immunofluorescence staining

HiPSCs and mammary gland organoids (MGOs) were briefly washed in PBS and fixed in 4% paraformaldehyde (pH 7.4) for 20–30 min at room temperature. Samples were washed three times in PBS, permeabilized with 0.1–0.2% Triton X-100 in PBS for 20 min, and blocked in 2% BSA in PBS for 1 h at room temperature. Whole-mount organoids were incubated with primary antibodies diluted in 0.1% BSA (Table S1) for 2 h at room temperature or overnight at 4 °C. Following three PBS washes, samples were incubated with fluorophore-conjugated secondary antibodies, counterstained with DAPI, and, where indicated, rhodamine–phalloidin for 45 min at room temperature protected from light. Samples were washed three times in PBS and imaged either directly in PBS or mounted in 80% glycerol in PBS for long-term storage at 4 °C. Negative controls included omission of primary antibodies or incubation with secondary antibodies alone. Imaging was performed using a Nikon Eclipse Ti2 confocal microscope, and images were processed and analysed using CellProfiler version 4.2.8.

### Primary Human Breast Tissue

Primary human breast tissue was obtained from a 22-year-old nulliparous woman undergoing breast surgery at Mediclinic, Belgium. The donor reported oral contraceptive use, and the tissue was collected during the follicular phase of the menstrual cycle. Tissue collection and use were conducted as part of a clinical study approved by the Ethics Committee Research UZ/KU Leuven (study S68111), following written informed consent from the donor.

The tissue was mechanically and enzymatically dissociated as previously described by Linnemann et al.^61^. The resulting primary cell suspension was seeded onto culture plates coated with 2% bovine skin gelatin (Type B, Sigma-Aldrich, G9391) and maintained in Mammary Epithelial Cell Growth Medium (MEGM™ BulletKit™, Lonza, CC-3150), supplemented with 3 µM Y-27632 dihydrochloride (Tocris Bioscience, 1254) for the first 5 days of culture. Cells were expanded by serial passaging and subsequently used for transplantation experiments at passage 3.

### Mice

All animal experiments were approved by the Animal Welfare Committee of KU Leuven and conducted in accordance with institutional and national guidelines (protocol number Onco-149/2024). Female *NOD-scid;IL2Rg^null^* (NSG) mice aged 3-3.5 weeks were used for transplantation experiments. Animals were bred in-house at KU Leuven or obtained from Charles River and maintained under specific pathogen-free conditions with a 12 h light/dark cycle, controlled temperature and humidity, and ad libitum access to food and water.

### Transplantation of mammary gland organoids into the mouse mammary fat pad

Mice were anaesthetized using isoflurane inhalation (1.5–2% isoflurane in air). Following skin sterilization with betadine and alcohol, a 2–3 mm incision was made between the fourth and fifth inguinal nipples. The endogenous mammary fat pad was surgically cleared from the nipple to the lymph node by cauterization of supplying blood vessels.

Day-14 cryopreserved hiPSC-derived mammary gland organoids and cryopreserved undifferentiated hiPSCs (negative control) of donor #1 were rapidly thawed and cultured for an additional 6 days to obtain day-20 organoids prior to transplantation. Primary human mammary epithelial cells (positive control) were prepared in parallel from reduction mammoplasty tissue. Day-20 organoids, undifferentiated hiPSCs, and primary mammary epithelial cells were dissociated into single cells using Accutase, washed, and resuspended at 1 × 10⁵ cells in 50 μL of BORN medium supplemented with 10% Matrigel per injection. Cell suspensions were injected into the cleared mammary fat pad using a TB syringe. Incisions were closed using wound clips and absorbable sutures, and mice were monitored post-operatively according to institutional guidelines.

After transplantation, mice were maintained for 10 weeks to allow engraftment and tissue development. At the experimental endpoint, mice were euthanized in accordance with institutional animal welfare guidelines. Transplanted mammary fat pads were harvested and processed for downstream analyses.

### Tissue processing, whole-mount imaging and cryosectioning

At the experimental endpoint, transplanted mammary fat pads were harvested and fixed in 4% paraformaldehyde (PFA) at 4 °C. Following fixation, tissues were washed in phosphate-buffered saline (PBS). A subset of transplanted mammary fat pads was processed as whole mounts for immunofluorescence imaging. Whole-mount tissues were stained according to the immunofluorescence protocol described above and imaged using confocal microscopy to assess overall tissue architecture, epithelial outgrowth, and marker expression within the intact graft.

For cryosection-based analyses, fixed tissues were cryoprotected in 30% (w/v) sucrose in PBS until fully equilibrated, embedded in Optimal Cutting Temperature (OCT) compound (Sakura Finetek), and frozen at −80 °C. Frozen tissue blocks were sectioned at a thickness of 8–10 μm using a Leica CM3050S cryostat. Sections were mounted onto Superfrost Plus microscope slides (Fisher Scientific), air-dried, and stored at −20 °C until immunofluorescence staining and imaging. Representative sections spanning the transplanted region were collected for downstream histological and immunofluorescence analyses.

### Image quantification

Quantitative image analysis was performed using Fiji (ImageJ, v2.x) and CellProfiler (v4.2.8). Confocal z-stacks were first processed in Fiji to generate maximum-intensity projections for downstream analysis. Images were subsequently imported into CellProfiler for automated segmentation and fluorescence quantification.

Nuclei were identified using the DAPI channel based on intensity and object shape characteristics. Cell boundaries were defined using either a fixed-distance expansion from the segmented nuclei (12-pixel radius) or the CellProfiler propagation algorithm, depending on marker localization and image quality. Segmentation results were visually inspected to ensure accurate identification of individual cells and exclusion of imaging artefacts.

Fluorescence intensity measurements were extracted from the corresponding cellular compartments for each marker. Mean fluorescence intensity, integrated intensity, and cell counts were quantified on a per-cell basis and aggregated across organoids within each biological replicate. Background fluorescence was determined from cell-free regions of the image and subtracted prior to downstream analysis. Quantitative measurements were exported from CellProfiler and analyzed using GraphPad Prism 10 and R (v4.5.1).

### ScRNA-seq

Organoids of donor #1 were collected for single-cell transcriptomic profiling. Two biological replicates were prepared. Replicate 1 included naïve MGO, MGO treated with 17β-oestradiol (E2; 10 µM), and MGO treated with E2 (10 µM) plus 4-hydroxytamoxifen (4OHT; 1 µM). Replicate 2 included the same three conditions, with additional exploratory perturbations using bisphenol A (BPA; 1 µM) and diethylstilbestrol (DES; 10 nM). Compound concentrations were selected based on the known differences in oestrogenic potency and prior experimental optimization; therefore, the concentrations differed between compounds. DES is much more potent at the oestrogen receptor than BPA, therefore, concentrations were not intended to be equimolar across treatments.

Organoids were mechanically disrupted and enzymatically dissociated with Accutase for 10 min at 37 °C, followed by gentle trituration and filtration through a 40 µm cell strainer. Viable single-cell suspensions with low doublet content were processed using the TempO-LINC platform (BioSpyder). Cells were washed, counted, and fixed according to the TempO-LINC fixation protocol using formaldehyde-containing fixation buffer at 4 °C. Fixed cells were quenched, supplemented with RNase inhibitor, mixed with freeze buffer, and cryopreserved at −80 °C before shipment on dry ice to the BioSpyder service facility.

Fixed and permeabilized cells were hybridized in situ with TempO-Seq whole-transcriptome detector oligo pools. Correctly hybridized adjacent detector oligos were ligated and subjected to combinatorial ligation-based cell barcoding. Barcoded libraries were PCR-amplified and sequenced on an Illumina platform using paired-end sequencing. Reads were demultiplexed using the TempO-LINC pipeline, aligned to species-specific TempO-LINC probe reference sequences using BWA v0.7.17, and gene count matrices were generated using featureCounts v2.0.1. Downstream analyses were performed using Seurat v5.5.0.

### Preprocessing and quality control of scRNA-seq data

Cells from each sample were filtered to remove low-quality cells and potential doublets. Cells with fewer than 100 detected genes or lower than 500 transcript counts were excluded from downstream analyses. Cells with higher than 30% ribosomal transcripts were also filtered as they represented low-quality or dying cells. Putative doublets were identified and removed using DoubletFinder (v2.0.6). TempO-LINC libraries do not target mitochondrial genes; therefore, mitochondrial transcript filtering was not applied. Following quality control, normalized count matrices were used for downstream dimensionality reduction, clustering and differential expression analyses.

### Clustering, annotation, and reference mapping

Clustering of the scRNA-seq data was performed by integrating all samples using canonical-correlation-analysis (CCA)-based integration implemented in Seurat (v5.5.0). Filtered gene matrices from each sample were normalized using the NormalizeData function. Highly variable genes were identified using FindVariableFeatures with the vst method, selecting 5,000 variable genes per sample. Integration anchors were identified using FindIntegrationAnchors with the following parameters: k.filter = 30, anchor.features = 3000 and k.score = 30. Integrated expression matrices were generated using IntegrateData.

The integrated dataset was subsequently scaled and centered using ScaleData, followed by principal component analysis. The first 20 principal components were used for unsupervised clustering using FindNeighbors and FindClusters at a resolution of 0.3. Dimensionality reduction for visualization was performed using Uniform Manifold Approximation and Projection (UMAP) with RunUMAP using the same principal components used for clustering.

Clusters were further evaluated for potential low-quality or artifactual populations using a multi-step curation strategy. First, cluster-level quality control metrics, including nCount_RNA, nFeature_RNA and ribosomal transcript content, were assessed to identify outlier populations. Second, the top differentially expressed genes for each cluster were examined to identify clusters dominated by ribosomal, mitochondrial, stress-associated or artifactual transcriptional programs. Third, canonical lineage markers were assessed to identify biologically coherent cell populations and potential off-target differentiation states.

Cluster identities were assigned using differential gene expression analysis performed with FindAllMarkers using the Wilcoxon rank-sum test. Marker genes were ranked by average log2 fold change and adjusted P value. Cluster annotations were further refined using canonical lineage markers, including *EPCAM*, *KRT8*, *KRT18*, *KRT6A* and *GABRP* for epithelial populations; *COL6A3*, *POSTN* and *PDGFRB* for stromal/perivascular populations; and *PECAM1*, *CLDN5* and *CDH5* for endothelial populations.

To further validate cluster identities, Reference mapping was performed using the Seurat FindTransferAnchors and TransferData workflow against a published single-cell transcriptomic atlas of normal human breast tissue reported by Kumar et al.^46^. Cell labels were transferred using the ‘broad-cell type’ annotation provided in the reference dataset, rather than the more granular ‘author-cell type annotations’, to enable robust assignment of major mammary cell lineages.. Predicted cell identities obtained through label transfer were compared with cluster-specific marker gene expression patterns and manually curated to ensure biological consistency. Final cluster annotations were assigned based on concordance between reference-based predictions, canonical lineage markers, and the top differentially expressed genes within each cluster.

### Cell-cycle scoring

Cell-cycle activity was assessed at the single-cell level using the CellCycleScoring function implemented in Seurat. Cells were scored for the expression of established S-phase and G2/M-phase gene sets provided with Seurat. Cell-cycle scores were calculated from normalized gene-expression values by comparing the average expression of genes associated with each cell-cycle phase with that of matched control gene sets. Each cell was assigned to G1, S or G2/M phase according to its relative S-phase and G2/M-phase scores. Cell-cycle scores and phase assignments were added to the single-cell metadata and used to examine differences in proliferative state across cell types and experimental conditions.

### Differential expression and pathway analysis

Differential gene expression analysis was performed using the Seurat framework. Pairwise comparisons were conducted between MGO and MGO_BPA, MGO and MGO_DES, and MGO_E2 and MGO_E2+4OHT conditions. Differentially expressed genes were identified using the Wilcoxon rank-sum test implemented in the FindMarkers function. Genes expressed in at least 5% of cells in either comparison group were included in the analysis.

To identify biological pathways associated with exposure-induced transcriptional changes, Gene Set Enrichment Analysis (GSEA) was performed using the Fast Gene Set Enrichment Analysis (FGSEA) algorithm. Genes were ranked according to average log2 fold-change values obtained from differential expression analysis. Enrichment testing was performed against the Hallmark gene set collection from the Molecular Signatures Database (MSigDB). Pathways containing between 5 and 500 genes were evaluated, and enrichment significance was assessed using 100,000 permutations. Normalized enrichment scores (NES) and Benjamini–Hochberg adjusted P values were used to identify significantly enriched pathways.

### Pseudotime trajectory analysis

Pseudotime analysis was performed using Monocle3 (v1.4.27) to investigate exposure-associated epithelial state remodelling. Trajectory inference was restricted to epithelial populations, including Basal Cells, LumSec-Basal, Luminal Sec, Luminal Sec-Myo, Luminal Sec-Major, Luminal HR-Major and Luminal HR-Active cells. Raw count matrices were converted to Monocle3 cell_data_set objects and analyzed within the Monocle3 framework. The UMAP embedding generated in Seurat was used for trajectory reconstruction and principal graph learning. Untreated MGO cells were specified as the trajectory root, and pseudotime values were assigned relative to this baseline state.

Differences in pseudotime distributions between exposure groups were assessed using Kruskal–Wallis tests followed by Benjamini–Hochberg corrected pairwise Wilcoxon rank-sum tests. Summary statistics, including median and mean pseudotime values, were calculated for each exposure condition.

Trajectory-associated genes were identified using the Monocle3 graph_test function with the learned principal graph as the neighbourhood structure. Genes were ranked according to Moran’s I statistic, which quantifies spatial autocorrelation of gene expression along the inferred trajectory. Genes exhibiting significant trajectory-associated expression patterns were defined using a false discovery rate threshold of q < 0.05.

To identify biological pathways associated with epithelial state progression, Gene Set Enrichment Analysis (GSEA) was performed using the fgsea package. Genes were ranked according to Moran’s I values and tested against the Hallmark gene set collection from the Molecular Signatures Database (MSigDB v2023.1). Gene sets containing between 5 and 500 genes were included, and enrichment significance was assessed using 100,000 permutations. Pathways with a Benjamini–Hochberg adjusted P value < 0.05 were considered significantly associated with epithelial state progression.

### Cell–cell communication analysis

Cell–cell communication analysis was performed using LIANA (v0.1.14) implementing the CellPhoneDB ligand–receptor resource on integrated single-cell RNA sequencing data generated from human induced pluripotent stem cell (hiPSC)-derived mammary gland organoids. Ligand–receptor interactions were inferred using the liana_wrap() function with the CellPhoneDB method. Significant interactions were identified using a permutation-based P-value threshold of <0.05. To reduce spurious interactions arising from lowly expressed genes, ligand and receptor genes were each required to be expressed in at least 10% of cells within the corresponding sender and receiver populations. Additional stringency was applied by retaining only interactions with a mean interaction score >0.5.

Integrin-mediated interactions were excluded from downstream analyses to reduce dominance of extracellular matrix adhesion programs and facilitate interpretation of signalling-associated interactions. Interaction significance was visualized as −log10(P value), and interaction strength was represented by the mean ligand–receptor interaction score.

Annotated cell populations were grouped into epithelial and stromal compartments based on cluster identity. Cell–cell communication networks were examined globally and subsequently stratified into epithelial–epithelial, epithelial-to-stromal, stromal-to-epithelial, and stromal–stromal interaction categories to resolve lineage-restricted and microenvironmental signalling programs. Bidirectional epithelial–stromal communication networks were visualized using ligand–receptor interaction dot plots and circos diagrams, with interactions filtered according to the significance and expression thresholds described above.

### Bulk transcriptomic analysis

Bulk transcriptomic count data were processed at the gene level prior to differential expression analysis. Low-depth samples were removed using a library-size threshold defined as the mean library size minus two standard deviations. Lowly detected probes were excluded using a relevance filter requiring detection in at least 75% of samples within at least one replicate group. Where multiple probes mapped to the same gene, counts were summed to generate a single gene-level count matrix.

Differential expression analysis was performed using DESeq2 (v1.50.2). Variance-stabilized counts generated using the varianceStabilizingTransformation function in DESeq2 were used for principal component analysis and sample-to-sample distance calculations. Differential expressions were modelled using a design formula including replicate and genotype effects (∼REPLICATE+CELL_LINE). Genes with adjusted P value < 0.05 and absolute log2 fold-change > 1 were considered differentially expressed.

Gene Ontology enrichment analysis was performed on significantly upregulated and downregulated genes using clusterProfiler (v 4.18.4) with human gene annotations from org.Hs.eg.db (v3.22.0). Enrichment results were adjusted for multiple testing using the Benjamini–Hochberg method.

Gene Set Enrichment Analysis was performed using the fgsea (v1.36.2) algorithm. Genes were ranked using the DESeq2 Wald statistic, and enrichment was tested against the Hallmark gene set collection from the Molecular Signatures Database. Gene sets containing between 5 and 500 genes were included. Normalized enrichment scores and Benjamini–Hochberg adjusted P values were used to identify enriched pathways.

Volcano plots were generated from DESeq2 results using EnhancedVolcano (v1.28.2). Significance thresholds were set at adjusted P value < 0.05 and absolute log2 fold-change > 1. Heatmaps of differentially expressed genes and curated pathway genes were generated from variance-stabilized expression values. Expression values were scaled by gene to generate row Z-scores, and heatmaps were visualized using ComplexHeatmap (v2.26.1).

### RNA isolation and quantitative PCR

Total RNA was isolated from organoids using the RNeasy Mini Kit (Qiagen) according to the manufacturer’s instructions. RNA concentration and purity were assessed using a NanoDrop spectrophotometer (Thermo Fisher Scientific). Complementary DNA (cDNA) was synthesized from total RNA using the iScript cDNA Synthesis Kit (Bio-Rad) following the manufacturer’s protocol.

Quantitative real-time PCR (qPCR) was performed using SYBR Green Master Mix (Applied Biosystems) on a QuantStudio 5 Flex Real-Time PCR System (Applied Biosystems). Gene-specific primers are provided in Table S1. Amplification was carried out under standard cycling conditions recommended by the manufacturer and melt curve analysis was performed to confirm amplification specificity.

Gene expression levels were normalized to the housekeeping gene GAPDH. Relative gene expression was calculated using the comparative Ct (2−ΔΔCt) method, with expression values reported relative to the indicated control condition. Each experiment was performed using independent biological replicates, and results are presented as mean ± standard error of the mean (SEM) unless otherwise stated. Graphs and statistical visualisations were generated using GraphPad Prism 10.

## Statistical analysis

All statistical analyses were performed in R (v4.5.1). Unless otherwise indicated, comparisons between two groups were performed using two-sided Wilcoxon rank-sum tests. Multiple-testing correction was performed using the Benjamini–Hochberg method. For pseudotime analyses, differences among exposure groups were assessed using Kruskal–Wallis tests followed by Benjamini–Hochberg corrected pairwise Wilcoxon rank-sum tests. Adjusted P values < 0.05 were considered statistically significant.

---

**Supplementary Figure 1.**
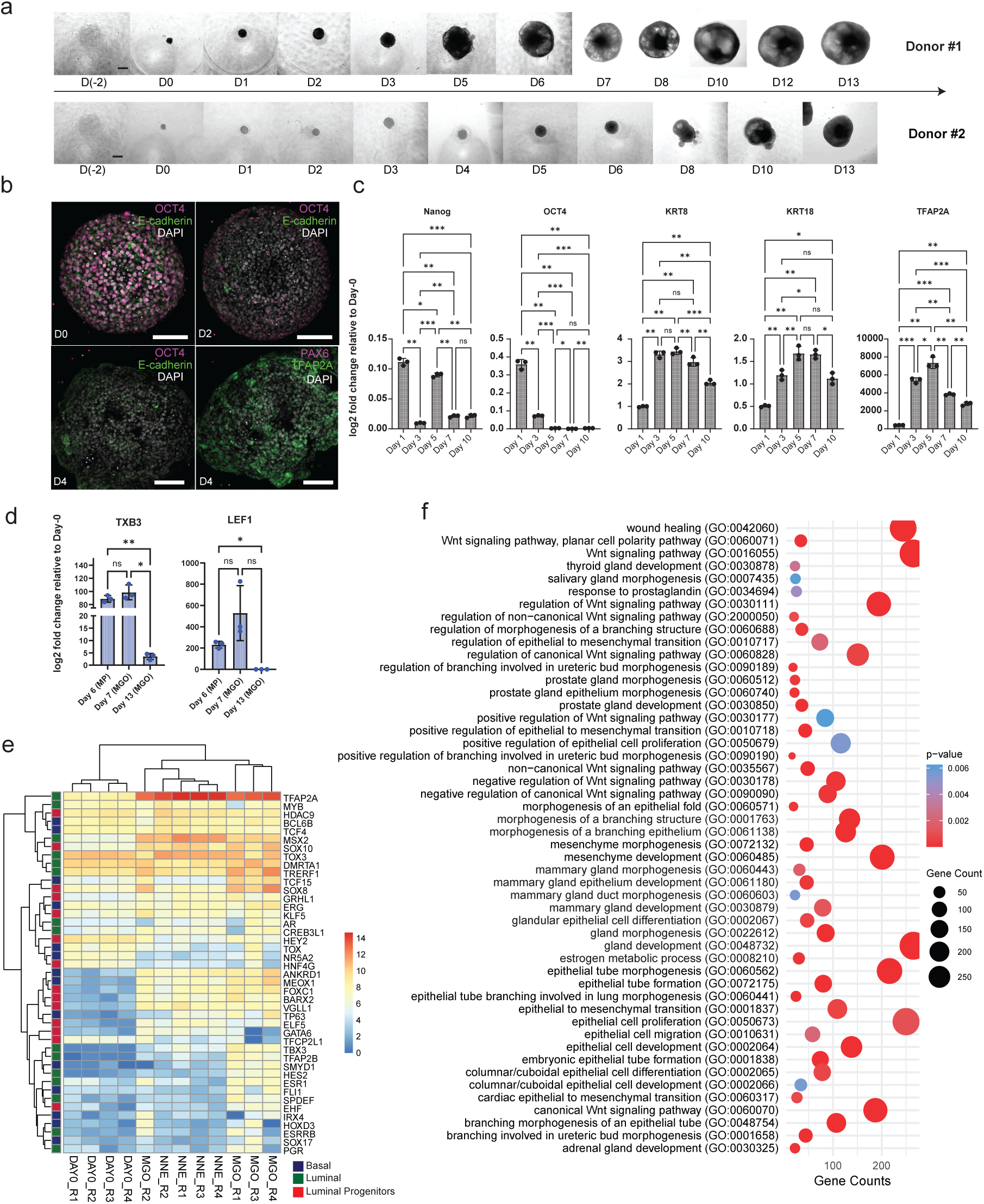
Optimization and validation of directed differentiation toward surface ectoderm and mammary fate. **a.** Brightfield imaging of morphological progression during directed differentiation. Representative brightfield images showing morphological progression of two independent hiPSC lines (donor #1 and donor #2)) during direct-ed differentiation from day −2 to day 13. Scale bars, 100 μm. **b.** Immunofluorescence at days 0, 2, and 4 for OCT4 (magenta) and E-cadherin (green). D4 spheroids were further stained for absence of PAX6 (magenta) and presence of TFAP2A (green. Nuclei were stained with DAPI (white). Scale bars, 100 μm. **c.** Quantitative RT–PCR analysis of pluripotency and ectoderm-associated gene expression during differentiation. Expression of *NANOG*, *OCT4, KRT8*, and *TFAP2A* was assessed across days 0–10. Mean ± s.d., where n = 3 biological replicates, each replicate having 8 spheroids. Significance assessed by one-way ANOVA with multiple comparisons. *p < 0.05, **p < 0.01, ***p < 0.001. **d.** Quantitative RT–PCR analysis of mammary placode-associated transcription factors *TBX3* and *LEF1* during placodal specification and organoid maturation across days 6–13. Mean ± s.d., n = 3. *p < 0.05; ns, not significant. Significance assessed by one-way ANOVA with multiple comparisons. **e.** Bulk RNA sequencing identifies stage-specific transcriptional programs during differentiation. Heatmap with log2-normalized expression of selected transcription factors associated with luminal (green), basal (blue), and luminal progenitor (red) populations across developmental stages and replicates (DAY0, NNE, and MGO). Rows represent transcription factors and columns represent individual samples. Gene expression is color-coded from low (blue) to high (red). Hierarchical clustering of both genes and samples reveals distinct lineage-associated transcriptional programs and stage-specific regulatory patterns within mammary epithelial differentiation. **f.** Gene Ontology enrichment analysis of differentially expressed genes during differentiation.

**Supplementary Figure 2.**
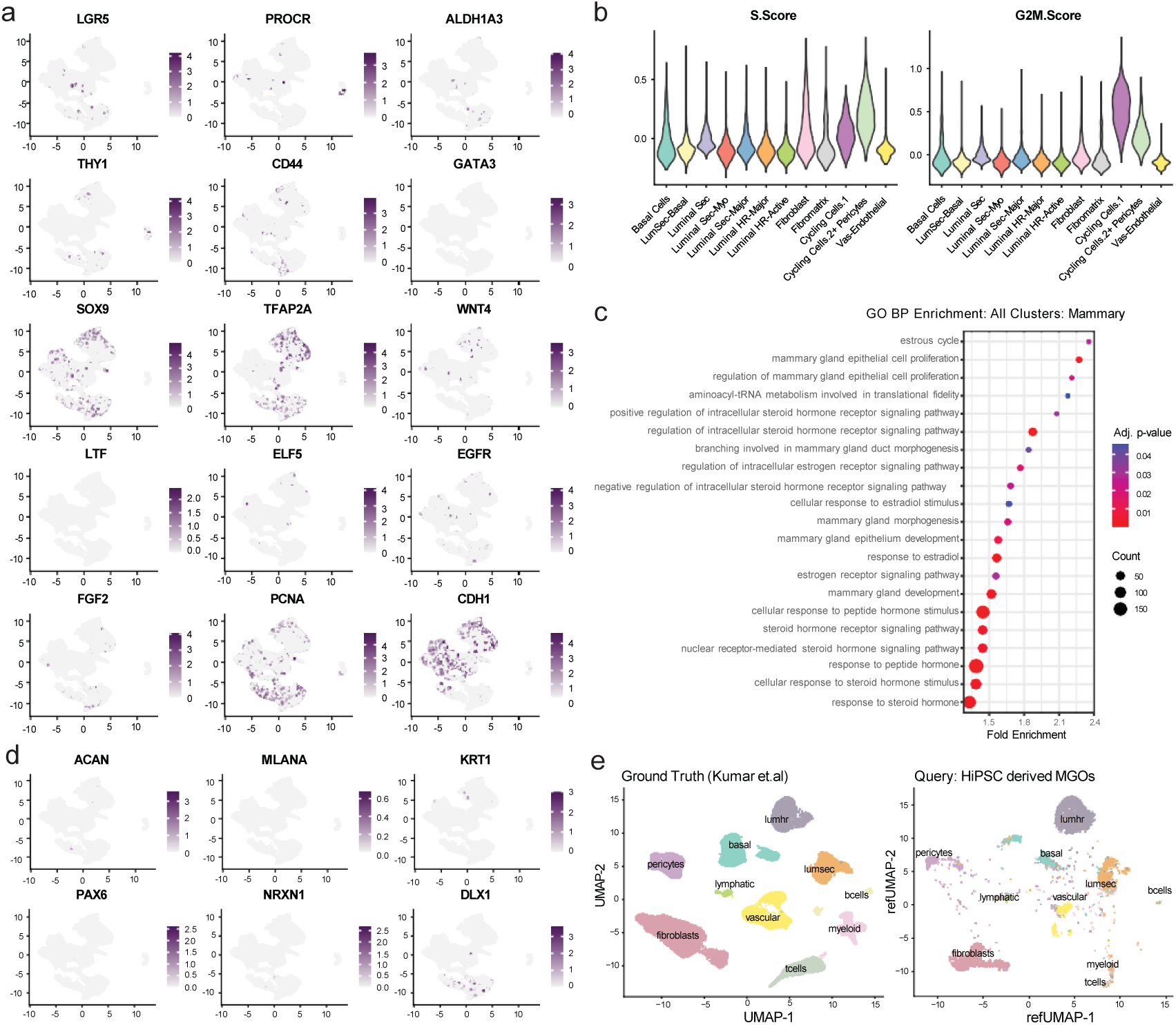
Single-cell validation of mammary epithelial identity and reference mapping. **a.** Single-cell expression of mammary epithelial lineage-associated and progenitor markers. UMAP single-cell RNA sequencing feature plots showing expression of additional representative mammary epithelial lineage and progenitor markers across integrated single-cell transcriptomes. Markers include stem/progenitor-associated genes (*LGR5*, *PROCR*, *ALDH1A3*, *THY1*, *CD44*, *SOX9*), luminal markers (*GATA3*, *ELF5*, *LTF*, *WNT4*), epithelial markers (*CDH1*, *EGFR*, *FGF2*, *PCNA*), and the non-neural ectoderm marker *TFAP2A* (n = 19020 cells pooled from two biological replicates, for each replicate n=10 organoids/condition). **b.** Cell-cycle activity across identified cell populations. Cell-cycle scoring across identified cell populations showing S-phase and G2/M-phase activity. **c.** Gene Ontology enrichment of cluster-associated differentially expressed genes. Gene Ontology (GO) enrichment analysis of differentially expressed genes across all clusters demonstrates enrichment of mammary gland development, mammary gland morphogenesis, oestrogen receptor signalling, steroid hormone signalling and mammary epithelial proliferation. **d.** Expression of non-mammary lineage-associated markers. UMAP feature plots demonstrating minimal expression of non-mammary lineage markers (*ACAN*, *MLANA*, *KRT1*, *PAX6*, *NRXN1* and *DLX1*) within the integrated single-cell transcriptomes. **e.** Reference mapping of hiPSC-derived mammary gland organoids to the human breast single-cell atlas. Reference mapping of hiPSC-derived mammary gland organoid cells to the human breast single-cell atlas described by Kumar et al. Ground-truth reference populations (left) and transferred annotations of organoid-derived cells (right) demonstrate transcriptional correspondence between MGOs and reference adult human mammary epithelial and stromal populations.

**Supplementary Figure 3.**
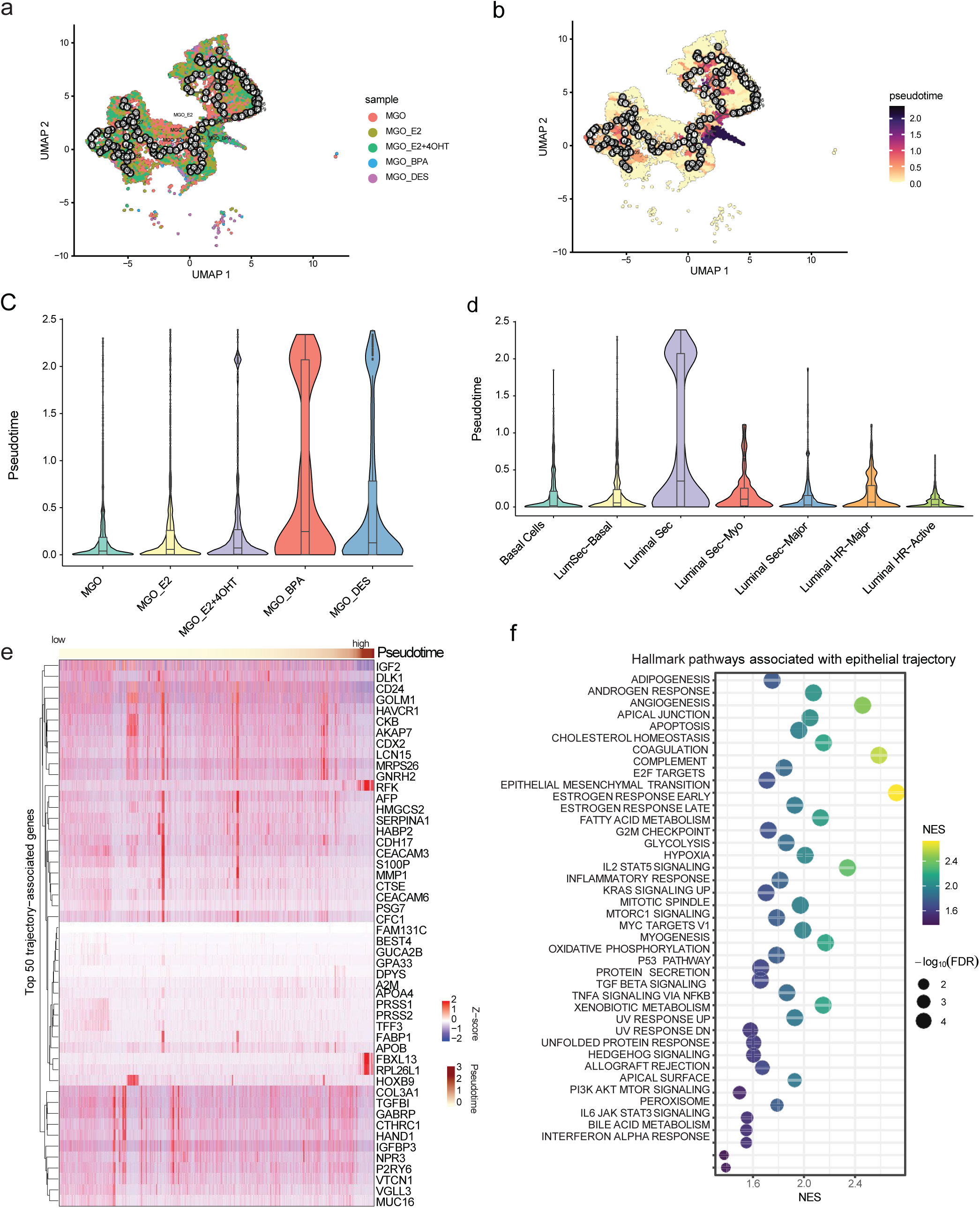
Endocrine perturbation remodels epithelial state trajectories. a. UMAP of integrated single-cell RNA-sequencing data showing epithelial cells from untreated (MGO), 10 µM E2-treated (MGO_E2), 10 µM E2 + 1 µM 4OHT-treated (MGO_E2+4OHT), 1 µM BPA-treated (MGO_BPA), and 10 nM DES-treated (MGO_DES) organoids following trajectory inference. Black circles and connecting lines indicate the inferred principal trajectory graph used for pseudotime estimation. Each coloured point represents a single epithelial cell, (n = 19,020 epithelial cells pooled from two biological replicates; 10 organoids/condition/replicate). b. Monocle3 pseudotime projected onto the epithelial trajectory. **c.** Violin plots of Monocle3 pseudotime values for individual epithelial cells, grouped by treatment condition **d.** Violin plots of Monocle3 pseudotime values for individual epithelial cells, grouped by epithelial cell subtype. **e.** Heatmap showing the expression patterns of the top 50 genes associated with the inferred epithelial trajectory, selected based on significant Moran’s I from the Monocle3 principal-graph test (q < 0.05, Moran’s I > 0.1). Columns represent individual epithelial cells ordered from low to high pseudotime (left to right), and rows represent trajectory-associated genes. Gene expression was z-score scaled across cells for each gene, with red indicating relatively higher and blue indicating relatively lower expression. The top annotation indicates the corresponding pseudotime value. **f.** Hallmark pathway enrichment associated with epithelial trajectory progression. Circles represent significantly enriched Hallmark gene sets identified from genes varying along the epithelial trajectory; circle colour indicates normalized enrichment score (NES), and circle size indicates statistical significance (−log10 adjusted P value).

**Supplementary Figure 4.**
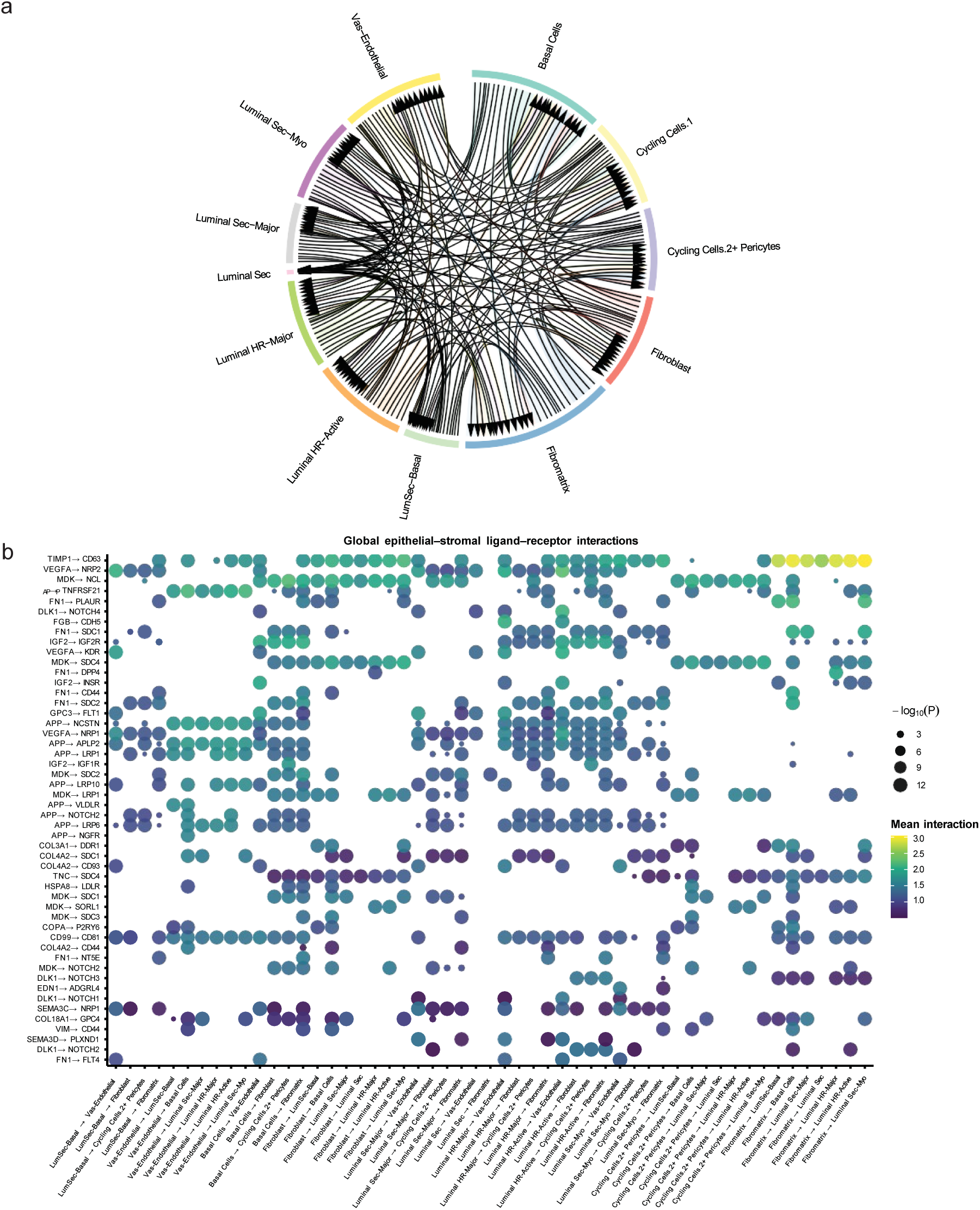
Cell–cell communication network within hiPSC-derived mammary gland organoids. **a.** Inferred ligand–receptor interaction network across cell populations. Cell–cell communication network inferred from integrated single-cell RNA-sequencing data using ligand–receptor interaction analysis. Global ligand–receptor interaction network inferred between epithelial and stromal cell populations. Chord plots illustrate predicted signalling interactions among basal, luminal, fibroblast, fibromatrix, endothelial and pericyte populations. **b.** Representative epithelial–stromal ligand–receptor interactions. Bubble plot showing representative epithelial–stromal ligand–receptor interactions ranked by interaction significance and mean interaction strength. Dot size represents −log10(P), and colour indicates the mean interaction score. Representative signalling pathways include extracellular matrix, growth factor, developmental, and angiogenic interactions.

